# Structural clustering of paired αβ T-cell receptor repertoires recovers antigen-specific convergence beyond sequence similarity

**DOI:** 10.64898/2026.11.22.768390

**Authors:** Devinder Srai, Dirk Aarts, John Frater, Matthew Raybould

## Abstract

T-cell receptor (TCR) antigen recognition depends on the three-dimensional geometry of the paired αβ complementarity-determining regions (CDRs), which primary sequence does not fully capture. Most repertoire-clustering methods group receptors by CDR3 motifs, sequence distance or V-J gene usage, and may miss receptors that differ in sequence but adopt similar loop conformations. We present SPACE-TCR (Structural Profiling of Alpha-Beta Clonotypes by Epitope), which stratifies predicted paired αβ backbone models by CDR length, aligns them on conserved framework anchors and clusters them by Cα RMSD across the CDRs. On an independently curated benchmark of 4,779 paired CD8 TCRs spanning 367 epitopes, 94.6% of SPACE-TCR clusters at the focal 0.75 Å threshold had purity ≥ 0.9, close to ClusTCR (97.3%), while clustering more than twice as many unique CDR3β (10.4% versus 4.7%); every method that clustered more did so at markedly lower purity. SPACE-TCR had the lowest polyspecificity (5.4%), and 22.7% of its high-purity clusters were sequence-diverse or sequence-unrelated, categories rarely recovered by the evaluated CDR3β-similarity methods. With increasing unannotated repertoire background, it kept higher purity than the evaluated sequence-based methods across all available comparisons (30.4% at 89.2% noise, compared with 15.0% for TCRMatch). Paired structural clustering therefore complements sequence-based analysis, recovering convergent receptor families that sequence criteria miss.

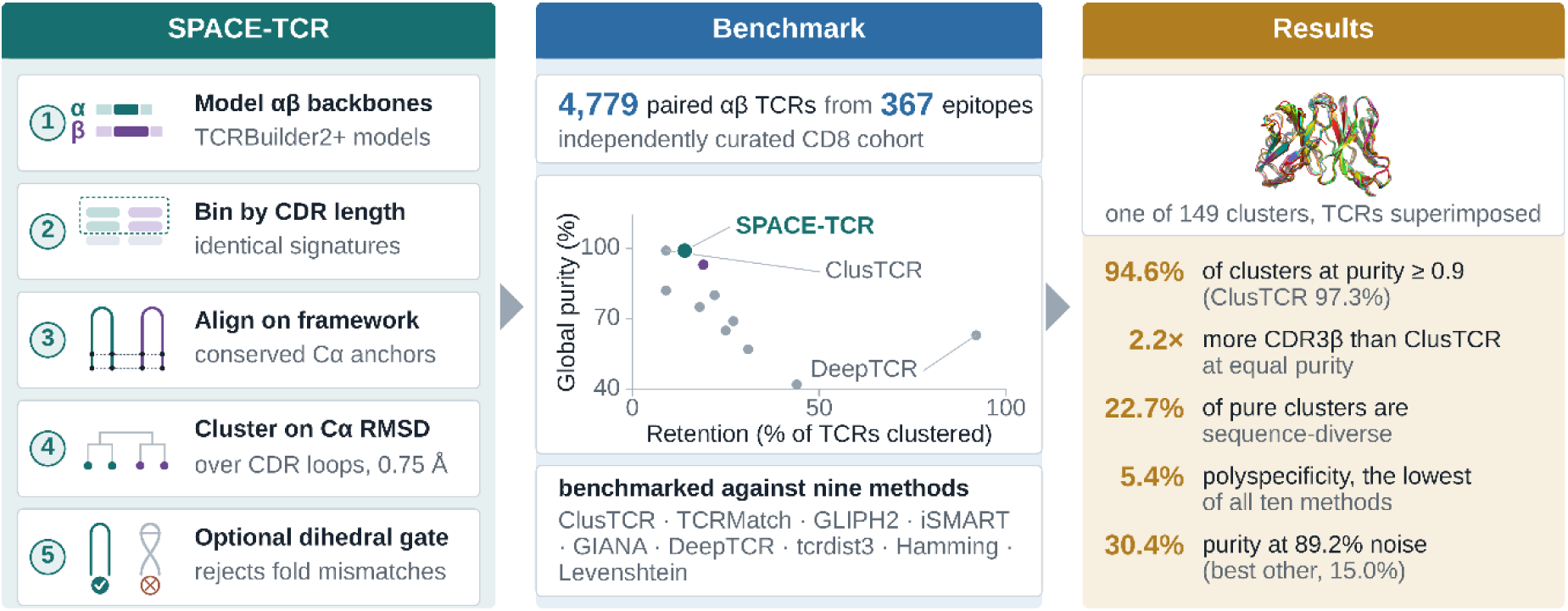

## 1. Introduction

T-cell receptor antigen recognition depends on the three-dimensional conformation of the complementarity-determining regions (CDRs), their arrangement across the paired αβ heterodimer, and the geometry of the resulting TCR-pMHC interface. Primary sequence does not provide these structural features directly. Considerable effort has gone into modelling the TCR-pMHC complex itself, and it remains a hard problem, because three things vary at once and interact: the receptor’s six CDR loops, the peptide, and the MHC allele presenting it. The difficulty is not only combinatorial but conformational. Comparing 391 bound and unbound structures across 22 TCRs, 19 MHC alleles and 79 peptides, McMaster et al. (2024) found that every CDR loop moves on engagement—by 2.38 Å on average for CDR3α and 1.50 Å for CDR3β— the CDR1 and CDR2 loops move as near-rigid bodies, while only the CDR3 loops deform intrinsically, and most of the change on the pMHC side falls in the peptide rather than the MHC. A recent evaluation of thirteen TCR–pMHC predictors reported that methods are correspondingly weakest at exactly the loops and the docking geometry that determine specificity, with docking orientation remaining the dominant error mode (Lu et al., 2026). Given these limitations, we use structural modelling to predict the unbound receptor alone, where prediction is comparatively accurate, and ask whether the relative geometry of its CDR loops is enough to group receptors that share a specificity.

The potential for identifying shared antigen specificity through receptor similarity was established by Glanville et al. (2017) and Dash et al. (2017), who demonstrated that TCRs recognising the same epitope can exhibit conserved sequence motifs and form distinguishable groups. Subsequent clustering methods have predominantly represented receptor similarity at the sequence level. ClusTCR, GLIPH2, TCRMatch, iSMART, GIANA and DeepTCR, alongside conventional Hamming and Levenshtein distance approaches, primarily exploit CDR3β sequence information, whereas tcrdist3 incorporates weighted distances across paired αβ sequences. These methods can efficiently identify clonotypes sharing close sequence homology or conserved motifs, but do not explicitly measure the three-dimensional conformations of the antigen-binding loops. Systematic comparison of them finds that no sequence-based model dominates, and that measured performance depends heavily on the epitope and on the evaluation design (Hudson et al., 2024). Consequently, receptors with divergent CDR3 sequences may adopt similar backbone geometries yet remain separated by sequence-based clustering. Conversely, similar CDR3 motifs may occupy distinct structural environments owing to differences in the paired chain, germline-encoded CDR regions or surrounding framework. Structural comparison may therefore capture relationships that are not readily apparent from sequence similarity alone.

The incorporation of predicted TCR structures into specificity analysis is increasingly established. The 2026 VDJdb release, for example, used TCRmodel2 to model paired-chain receptor records and assess them according to structural prediction confidence (Luppov et al., 2026). In parallel, representation-learning approaches have been applied to the same problem, although contrastive and masked-language models have so far struggled to outperform sequence-alignment baselines (Nagano et al., 2025). These uses of structure are, however, annotation rather than grouping: predicted complexes are used to score the plausibility of individual receptor–epitope records, while the clustering itself remains sequence-based, whether by motif embeddings or by edit distance. Identifying functionally related TCRs beyond sequence similarity has important implications for repertoire analysis. In particular, grouping structurally convergent receptors could enable the identification of shared antigen-specific responses across individuals, the detection of disease-associated receptor families, and the tracking of immune responses following infection, vaccination or therapeutic intervention. Such approaches may be particularly valuable in largely unannotated clinical repertoires, where individual clonotypes are frequently private and the antigen specificity of most receptors remains unknown.

Here, we ask the complementary question of whether predicted geometry can serve as the clustering metric itself by adapting the antibody-clustering framework SPACE2 (Spoendlin et al., 2023) to paired αβ TCRs. SPACE-TCR (Structural Profiling of Alpha-Beta Clonotypes by Epitope) aligns receptors using conserved framework residues before calculating Cα root-mean-square deviation (RMSD) across the CDR loops, thereby quantifying differences in antigen-binding loop geometry independently of overall model orientation. By preserving native αβ pairing, SPACE-TCR captures the combined structural contribution of both receptor chains rather than relying on individual CDR3 sequences. We assess whether this geometry-based representation identifies epitope-associated receptor clusters, and benchmark it against the nine sequence-based methods evaluated by Jouannet et al. (2025) on their independently curated, epitope-annotated TCR cohort. We then apply the same comparison to their noise benchmark, in which the annotated receptors are held fixed while unlabelled background sequences are progressively added. To our knowledge, this is the first structure-based method for T-cell receptor clustering and the first systematic benchmark of structural clustering performance in both annotated reference databases and the presence of repertoire background noise.

## 2. Methods

### 2.1 Structural modelling and confidence estimation

Where experimental records did not contain full-length sequences, germline V-gene segments were reconstructed with Stitchr to standardise framework regions and CDR boundaries before modelling (Heather et al., 2022). Full-length paired sequences were then modelled with TCRBuilder2+ (Quast et al., 2025; using OpenMM 8.2) and exported as PDB files. We retained all successfully modelled receptors.

The analysis is intended to preserve relative loop geometry and structural neighbourhoods rather than exact coordinates for docking or energy calculations. SPACE-TCR therefore assumes that receptors modelled with the same predictor and workflow retain sufficiently consistent relative geometry for neighbourhood analysis, even when the absolute coordinates of an individual loop are uncertain. All models here are unbound (apo) predictions. As CDR3 loops move by 1.5-2.4 Å on engagement (McMaster et al., 2024), which exceeds the clustering threshold evaluated here, SPACE-TCR does not claim that two receptors within 0.75 Å adopt the same bound conformation; it claims that their predicted apo geometries are near-identical, and tests empirically whether that predicts shared specificity. The thresholds reported here are properties of this model ensemble and would require recalibration for another predictor.

### 2.2 The SPACE-TCR clustering algorithm

SPACE-TCR has three stages. Predicted models are first divided into bins with identical CDR-length signatures, so that RMSD is calculated only between geometrically comparable loops. Structures within each bin are then aligned on Cα atoms. Finally, Cα RMSD is calculated across the CDR residues, and receptors are grouped at a fixed structural threshold, set in ångström. An exhaustive all-pairs comparison is prohibitive at repertoire scale. SPACE2 implementation was modified to allow moderate-sized bins to be clustered exactly by constructing an RMSD-threshold graph and extracting connected components. Large bins use a greedy representative procedure: each model is assigned to the first representative within threshold or becomes a new representative. This hybrid approach preserves exact near-neighbour clustering where feasible and permits analysis of cohorts containing hundreds of thousands of models. All results reported here were produced with the exact connected-components procedure. Additionally, a dihedral distance output was added to the original SPACE2 implementation, enabling further pruning post-distance-based clustering to remove those TCRs that cluster based on RMSD but differ in topology. RMSD measures Cα superposition, whereas φ and ψ angles describe local backbone torsion; the gate therefore excludes loops with similar Cα positions but different internal folds. Clustering immune-receptor loops by backbone geometry has an antibody precedent: North et al. (2011) classified antibody CDR conformations into canonical clusters from their φ/ψ dihedrals, and the same distinction between position and torsion motivates the gate used here.

### 2.3 Benchmark cohort and epitopes

The epitope-level benchmark used the curated CD8 cohort of Jouannet et al. (2025). As per Jouannet, records were only retained when Verified = 2, Identification > 4.3, both CDR3α and CDR3β were present, all V- and J-gene calls were available, and the epitope had at least two unique paired entries. TCRBuilder2+ successfully modelled 4,031 of the 4,779 paired TCRs across 367 epitopes. All 4,779 formed the SPACE-TCR input, so retention is measured against the full cohort and the 748 receptors without a model count as unclustered. As per the original benchmark, we highlight six epitopes in this work, spanning common and rare epitopes in this dataset: GILGFVFTL (influenza A M1), NLVPMVATV (CMV pp65), GLCTLVAML (EBV BMLF1), FLCMKALLL (HIV Gag), NYNYLYRLF (HIV Nef), and LLFGYPVYV (HTLV-1 Tax). We evaluated RMSD thresholds of 0.25, 0.50, 0.75, 1.00, and 1.25 Å, both without a dihedral gate and with a dihedral angle gate across the CDR3 regions of both chains. Unless otherwise stated this angle was set at a threshold of 0.5-radian or 28 degrees. The 0.75 Å RMSD setting was designated as the focal operating point to balance high purity and relatively high retention. More stringent thresholds can be implemented by users where purity is favoured over retention. For direct comparison, SPACE-TCR distances were evaluated with the Jouannet purity, retention, and sensitivity definitions and compared with ClusTCR (Valkiers et al., 2021), GLIPH2 (Huang et al., 2020), TCRMatch (Chronister et al., 2021), iSMART (Zhang et al., 2020), GIANA (Zhang et al., 2021), DeepTCR (Sidhom et al., 2021), tcrdist3 (Mayer-Blackwell et al., 2021), and Hamming- and Levenshtein-distance baselines (Hamming, 1950; Levenshtein, 1966).

The Jouannet benchmark pools IEDB (Vita et al., 2025), VDJdb (Luppov et al., 2026) and McPAS-TCR (Tickotsky et al., 2017); the epitope labels therefore originate with those resources. The focal value is lower than the 1 Å used for loop-level RMSD clustering of TCRBuilder2+ predictions by Quast et al. (2025), and the difference is deliberate rather than incidental. That work clusters to measure structural diversity, where a looser threshold gives a more informative partition of the space; here clusters are proposed as specificity groups, and the operating point is chosen on labelled data against the purity-retention trade-off rather than on the geometry alone. At 1 Å purity falls while retention rises (Table 1), and 0.75 Å is where purity is still at the high-purity end of the published methods. Both values are reported throughout so the two conventions can be compared directly. The cohort is small beside the raw size of the public databases, which now list over 200,000 TCR-epitope records (Luppov et al., 2026), but the informative comparison is with paired, confidently annotated, class I-restricted receptors. Computed from the 2026 VDJdb release, that subset holds 3,316 paired receptors at confidence ≥ 1 and 1,616 at the highest confidence tier, against 4,779 here; the headline totals are dominated by unpaired β-only records carrying no confidence score. The cohort is therefore large relative to publicly available datasets of paired, confidently annotated, class I-restricted TCRs.

**Table 1.** SPACE-TCR performance across five RMSD thresholds, with RMSD-only and dihedral-gated modes scored using the Jouannet et al. (2025) definitions. The focal 0.75 Å RMSD-only setting is highlighted.

| Threshold (Å) | Mode | n clusters | TCRs clustered | Retention | Global purity | % clusters purity ≥ 0.9 |
|---|---|---|---|---|---|---|
| 0.25 | RMSD | 21 | 65 | 1.4% | 100.0% | 100.0% |
| 0.50 | RMSD | 95 | 393 | 8.2% | 99.0% | 95.8% |
| **0.75** | **RMSD (focal)** | **149** | **681** | **14.2%** | **98.7%** | **94.6%** |
| 0.75 | RMSD+dih | 139 | 658 | 13.8% | 98.6% | 94.2% |
| 1.00 | RMSD | 202 | 1,000 | 20.9% | 92.4% | 75.2% |
| 1.00 | RMSD+dih | 172 | 903 | 18.9% | 92.7% | 76.2% |
| 1.25 | RMSD | 321 | 1,555 | 32.5% | 78.4% | 41.7% |
| 1.25 | RMSD+dih | 221 | 1,186 | 24.8% | 84.3% | 53.8% |

### 2.4 Evaluation metrics

Metrics follow the Jouannet implementation; the formal definitions are given in Supplementary Section S18 (Eqs S1–S10). Every non-singleton cluster has a dominant epitope: the label most of its members carry. The receptors that carry it are the ones the cluster has grouped correctly, and the two purity measures differ only in what they average over. Global purity pools every clustered receptor and asks what fraction sit with their own cluster’s dominant label, so a large cluster influences it more than a small one. Per-cluster purity asks the same question inside one cluster, and a cluster is high-purity when the answer reaches 0.9; counting those clusters gives the percentage of clusters at purity ≥ 0.9, the column Tables 1 and 2 are ordered on, and counting their members gives the percentage of clustered TCRs in high-purity clusters. A method can therefore score well on one and less well on the other: at the focal threshold a few small mixed clusters hold the cluster proportion to 94.6% while global purity, which they barely move, stands at 98.7%. Retention is the fraction of all input TCRs placed in a non-singleton cluster. The methods evaluated take different inputs (e.g. paired receptors, CDR3β with its gene calls), so every cross-method comparison counts the same unit: the cohort’s 4,292 unique CDR3β sequences.

**Table 2.** Comparison of SPACE-TCR with the nine sequence-based methods evaluated by Jouannet et al. (2025). Published methods are ordered by the % purity ≥ 0.9 column, the proportion of non-singleton clusters whose per-cluster purity is at least 0.9 (Eq. S9). Global purity (Eq. S1) is the fraction of all clustered sequences carrying their own cluster’s dominant epitope. The two answer different questions — one counts clusters, the other counts sequences — and rank the methods differently: DeepTCR reaches higher global purity than Levenshtein distance while forming a lower proportion of clusters with purity ≥ 0.9. The Signal column records what each method was given as input. CDR3β clustered is counted as in Section 2.4. Supplementary Table S1b lists exact output counts.

| Method | Signal | n clusters | CDR3β clustered (%) | Global purity | % purity ≥ 0.9 | Median LD | % polyspecific |
|---|---|---|---|---|---|---|---|
| **SPACE-TCR (RMSD, 0.75 Å)** | **paired CDR3αβ Cα backbone** | **149** | **446 (10.4%)** | **98.7%** | **94.6%** | **2.0** | **5.4%** |
| **SPACE-TCR (RMSD+dih, 1.0 Å)** | **paired CDR3αβ Cα backbone + dihedral** | **172** | **642 (15.0%)** | **92.7%** | **76.2%** | **3.0** | **26.2%** |
| ClusTCR | paired CDR3αβ sequence | 111 | 201 (4.7%) | 99.3% | 97.3% | 1.0 | 6.3% |
| TCRMatch | CDR3β sequence | 285 | 270 (6.3%) | 82.4% | 63.5% | 1.0 | 40.0% |
| GLIPH2 | CDR3β motif (paired input) | 380 | 713 (16.6%) | 79.8% | 51.0% | 1.0 | 30.5% |
| iSMART | CDR3β motif | 465 | 587 (13.7%) | 74.8% | 45.0% | 1.0 | 58.5% |
| Hamming distance | CDR3β sequence | 548 | 806 (18.8%) | 69.4% | 36.3% | 1.0 | 66.1% |
| GIANA | CDR3β sequence | 593 | 591 (13.8%) | 64.9% | 35.6% | 2.0 | 66.3% |
| Levenshtein distance | CDR3β sequence | 494 | 913 (21.3%) | 57.3% | 33.8% | 1.0 | 67.4% |
| DeepTCR | paired CDR3αβ + V/J (deep) | 1,628 | 3,920 (91.3%) | 63.2% | 19.7% | 6.0 | 80.9% |
| TCRdist3 | paired CDR3αβ distance | 669 | 1,940 (45.2%) | 42.2% | 7.0% | 9.0 | 92.7% |

Epitope-level sensitivity applies a stricter rule. A cluster qualifies for epitope e if it contains at least four TCRs, is dominated by e, and has at least 67% of its members labelled e; sensitivity is the fraction of all input TCRs labelled e that such clusters recover. The numerator counts distinct receptors, so a receptor assigned to several qualifying clusters is counted once. Epitope totals differ between the paired and CDR3β-keyed methods and are given in Supplementary Section S13. Polyspecificity is the fraction of non-singleton clusters containing more than one epitope label; it is stricter than purity, since a cluster that is 95% one epitope and 5% another remains high-purity but counts as polyspecific. TRBV and TRBJ composition use the same dominance calculation but are described as homogeneity, reported as the percentage of non-singleton clusters in which one gene accounts for at least 90% of members, with gene calls keyed to CDR3β. Within-cluster sequence diversity is the median pairwise Levenshtein distance among member CDR3 sequences, measured separately for each chain; CDR3α diversity is assessed only for methods with paired-chain cluster output.

### 2.5 Consensus clustering

To test whether combining structural and sequence signals improves on the purity of sequence-only methods, we formed consensus clusters that keep only the groupings both representations agree on, by intersecting SPACE-TCR assignments with sequence-based assignments. Two receptors were placed together only when both methods grouped them. Four combinations were evaluated: each of the two SPACE-TCR modes (0.75 Å RMSD-only and 1.0 Å RMSD+dih) with ClusTCR or GLIPH2.

### 2.6 Performance under high noise settings

The curated benchmark assigns an epitope label to every receptor, whereas most patient repertoires are largely unannotated. We therefore repeat the Jouannet et al. (2025) noise analysis using the same 10X Genomics dextramer dataset. A fixed set of 2,876 dextramer-annotated CDR3β sequences is progressively diluted with unannotated ‘noise’ TCRs from the same donor across six subsets containing 0, 62.2, 76.7, 83.2, 86.8, and 89.2% unannotated TCRs, or noise.

SPACE-TCR and the published outputs were scored on the same subsets, labels, purity definition, and minimum cluster size. This produced evaluation subsets of 2,478 to 22,560 sequences, with 2,374 antigen-specific, epitope annotated sequences in every subset. Purity and retention were assessed across these subsets. Only paired receptor sequences were included in this analysis as a control measure to ensure identical inputs between SPACE-TCR input and sequence-based methods.

## 3. Results

### 3.1 Threshold sweep and focal operating point

Table 1 summarises performance across the five RMSD thresholds. The focal 0.75 Å setting achieved 98.7% global purity at 14.2% retention, and 94.6% of its clusters had per-cluster purity ≥ 0.9. The tightest 0.25 Å setting reached 100% purity but retained only 1.4% of the input TCRs. Tightening the focal setting to 0.50 Å increased global purity above 99% while approximately halving retention. Loosening the threshold to 1.00 or 1.25 Å increased retention, with the 1.00 Å setting still remaining above 90% global purity. At 0.75 Å, the dihedral-gated mode gave nearly identical purity (98.6% versus 98.7%) and clustered slightly fewer TCRs (658 versus 681). The 0.75 Å RMSD-only setting was retained as the focal point because it lies near the inflection of the purity-retention curve (Figure 1).

**Figure 1.**
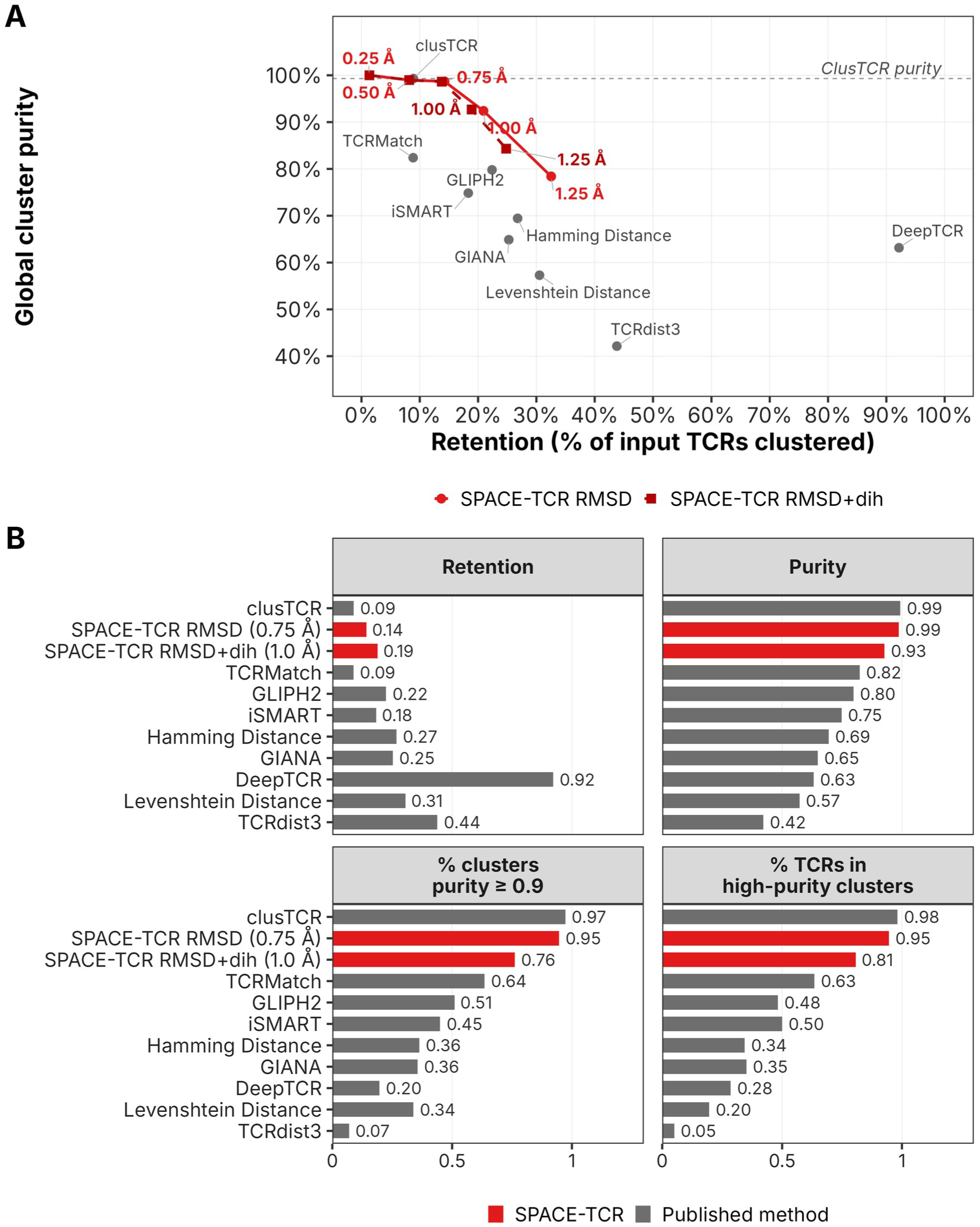
Benchmark performance on the curated CD8 cohort. (A) Purity-retention positions for the SPACE-TCR threshold sweep and nine published sequence-based methods. Circles show RMSD-only thresholds; squares show selected dihedral-gated settings. (B) Retention, global purity, proportion of clusters at purity ≥ 0.9, and proportion of TCRs in high-purity clusters. SPACE-TCR combines high purity with greater yield than the only comparably pure published method. Retention is counted as in Section 2.4. The Signal column of Table 2 records what each method was given as input.

### 3.2 Head-to-head comparison with sequence-based tools at the focal threshold

Figure 1A compares the SPACE-TCR threshold sweep with nine published methods. At 0.75 Å, SPACE-TCR achieved purity comparable to that of ClusTCR while retaining about twice as many receptors. Dihedral gating substantially improved clustering at looser RMSD thresholds. At 1.25 Å, a 0.5-radian gate increased global purity from 78.4% to 84.3%, high-purity clusters from 41.7% to 53.8%, and TCRs in high-purity clusters from 52.1% to 64.9%, while reducing retention by 7.7 percentage points. We therefore use RMSD-only at 0.75 Å for the primary comparison and reserve dihedral gating for looser thresholds.

Table 2 compares the focal 0.75 Å RMSD-only setting and the 1.0 Å dihedral-gated setting with the nine sequence-based methods. At the focal point, 94.6% of SPACE-TCR clusters had purity ≥ 0.9. Only ClusTCR was higher (97.3%); the next methods were TCRMatch (63.5%), GLIPH2 (51.0%), and iSMART (45.0%), with all remaining methods below 37%. SPACE-TCR clustered 446 unique CDR3β in 149 clusters against ClusTCR’s 201 in 111 — 2.2 times as many sequences at comparable purity. Every method that clustered more did so at lower purity: TCRMatch 6.3% at 63.5% high-purity clusters, GLIPH2 16.6% at 51.0%, TCRdist3 45.2% at 7.0% and DeepTCR 91.3% at 19.7%. SPACE-TCR also had the lowest polyspecificity (5.4%) and a higher median within-cluster CDR3β Levenshtein distance than the sequence methods (2 versus 1).

Figure 1B presents the four principal benchmark metrics together. SPACE-TCR combines high global purity, a high proportion of clusters above the 0.9 purity threshold, and a high proportion of TCRs within those clusters. Its retention is modest but exceeds that of ClusTCR, the only method with comparable cluster purity. DeepTCR has the highest retention but low purity.

### 3.3 Cluster size distribution

Cluster size affects the strength of downstream interpretation: two-member clusters provide limited support, whereas larger clusters permit more reliable assessment of heterogeneity and disease association. At 0.75 Å, 55.0% of SPACE-TCR clusters were pairs. Only GLIPH2 had a smaller share of pairs (38.2%), and it did so while reaching purity ≥ 0.9 in half its clusters against SPACE-TCR’s 94.6%. 25.5% contained at least five members, the highest proportion among high-purity methods — ClusTCR, the only method with higher cluster purity, reached five members in 12.6% of its clusters (Figure 2A). Mean cluster size was 4.6, compared with 3.8 for ClusTCR and 2.6 for TCRMatch. GLIPH2 (7.7) and the Hamming and Levenshtein baselines (4.1 and 5.2) produced larger mean clusters, but at much lower purity. Increasing the minimum cluster-size threshold left SPACE-TCR purity close to 99% from size 2 through size 20, indicating that its performance was not driven only by small clusters (Figure 2B).

**Figure 2.**
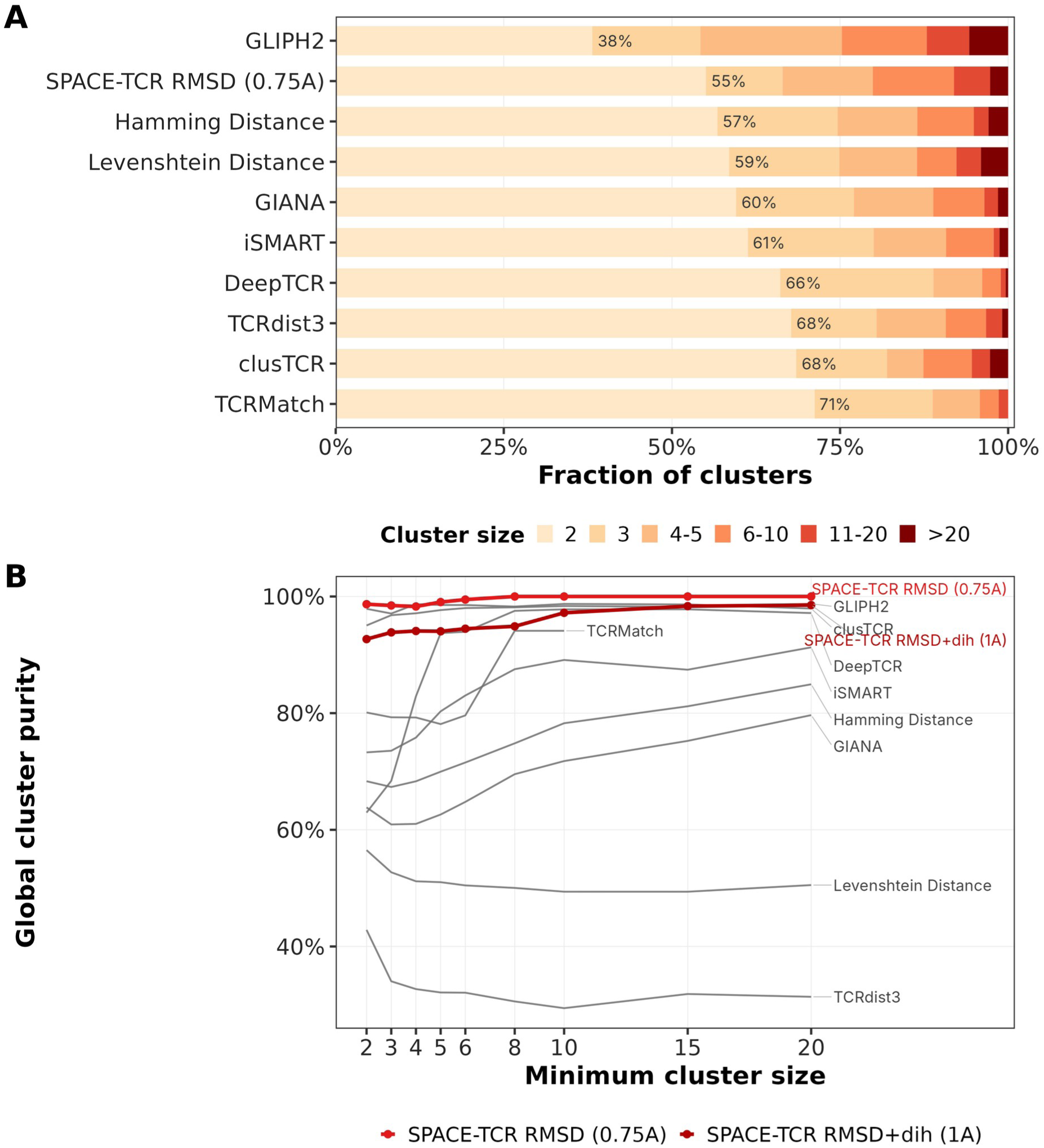
Cluster-size behaviour. (A) Distribution across methods. (B) Global purity (Eq. S1), not the proportion of clusters above 0.9 reported in Table 2, after progressively increasing the minimum cluster size from 2 to 20 members. Focal SPACE -TCR begins at 98.7% and reaches 100%; ClusTCR and GLIPH2 are the only published methods that stay above 95% throughout, and the edit-distance baselines do not rise above 85% at any threshold.

All cross-method comparisons used the same receptor set. SPACE-TCR received all 4,779 paired TCRs; TCRBuilder2+ produced models for 4,031 of them, and the remaining 748 count as unclustered. For GILGFVFTL, NLVPMVATV, and GLCTLVAML, the modelled input was a complete subset of the Jouannet cohort (454, 345, and 170 unique CDR3β, respectively, counted as in Supplementary Section S1), with more than 98.8% label agreement for each epitope. Comparisons for these three dominant specificities are therefore direct. FLCMKALLL and NYNYLYRLF were sparsely represented (81 and 46 unique CDR3β), and the focal mode had zero sensitivity for both. At these sizes few clusters can meet the qualifying rule of at least four members and 67% dominance, so zero sensitivity reflects that rule at small n rather than evidence that these receptors were incorrectly clustered.

### 3.4 Convergence classes: where structural clustering adds value

To determine where structural clustering added value, each high-purity focal SPACE-TCR cluster (purity ≥ 0.9, size ≥ 2; n = 141) was classified by its median pairwise CDR3β Levenshtein distance (Table 3). Of these clusters, 63 (44.7%) were sequence-identical or near-identical (LD ≤ 1), and 46 (32.6%) were sequence-similar (LD 2-3). A further 26 (18.4%) were sequence-diverse (LD 4-6), and six (4.3%) were sequence-unrelated (LD > 6). The latter two classes contain receptors with substantial CDR3β sequence differences but closely aligned backbone geometries. Together, 32 of 141 high-purity clusters (22.7%) fell outside the usual operating range of the benchmarked sequence methods. At 1.0 Å, this proportion increased to 61 of 152 high-purity clusters (40.1%), while global purity decreased only marginally to 92.4%. The same classification applied to every other method shows that sequence-divergent high-purity clusters are largely unique to the structural representation: the share is 0.0% for every edit-distance, motif and k-mer method evaluated and 1.4% for GLIPH2, against 22.7% here (Figure 4C). DeepTCR (48.3%) and tcrdist3 (71.4%) do form them, but by clustering loosely enough that only 19% and 7% of their clusters respectively reach purity ≥ 0.9.

**Table 3.** CDR3β sequence-divergence classes among high-purity SPACE-TCR clusters at 0.75 Å. Classes are defined by median within-cluster Levenshtein distance. The sequence-diverse and sequence-unrelated classes fall outside the usual operating range of the benchmarked sequence methods despite meeting the same purity threshold.

| **CDR3β sequence divergence class** | **Median CDR3β LD** | **Clusters (n)** | **Share of high-purity clusters** | **Recoverable by CDR3β-similarity methods?** |
|---|---|---|---|---|
| Sequence-identical / near-identical | ≤ 1 | 63 | 44.7% | Yes (ClusTCR, LD, HD) |
| Sequence-similar | 2–3 | 46 | 32.6% | Yes (GLIPH2 motifs) |
| Sequence-diverse | 4–6 | 26 | 18.4% | No |
| Sequence-unrelated | > 6 | 6 | 4.3% | No |
| **Total** | **—** | **141** | **100%** | **—** |

### 3.5 Per-epitope sensitivity

Epitope-level sensitivity varied across the six (out of 367) highlighted epitopes (Figure 3A). Values are computed as Eq. S4 specifies, over all input receptors carrying the epitope. Sensitivity as a metric combines recovery with a purity floor: a cluster contributes only if it holds at least four receptors and its dominant epitope covers at least 67% of them. Among clusters of four or more, this threshold discards 96% of the receptors tcrdist3 groups, 64% for Levenshtein, 50% for GIANA and 38% for Hamming, against 5% for SPACE-TCR under the dihedral gate and 1% at the focal threshold. Aggressive clustering therefore raises sensitivity only while the clusters stay above the purity floor. GILGFVFTL carries the most receptors and is therefore the epitope where the ranking is most informative (705 paired, 455 unique CDR3β); DeepTCR was highest at 0.413, with SPACE-TCR under the dihedral gate effectively level at 0.407, followed by GLIPH2 (0.367), the Hamming baseline (0.358), SPACE-TCR in RMSD-only mode (0.339), GIANA (0.310), iSMART (0.302), Levenshtein (0.278), ClusTCR (0.141), TCRMatch (0.096) and tcrdist3 (0.044). Averaged over the six epitopes, DeepTCR was highest (0.340), followed by SPACE-TCR with the gate (0.307) and in RMSD-only mode (0.265). Three of the six epitopes carry fewer than 90 receptors and FLCMKALLL yielded no qualifying cluster for any method, so per-epitope values at small n should not be read as ranking evidence.

**Figure 3.**
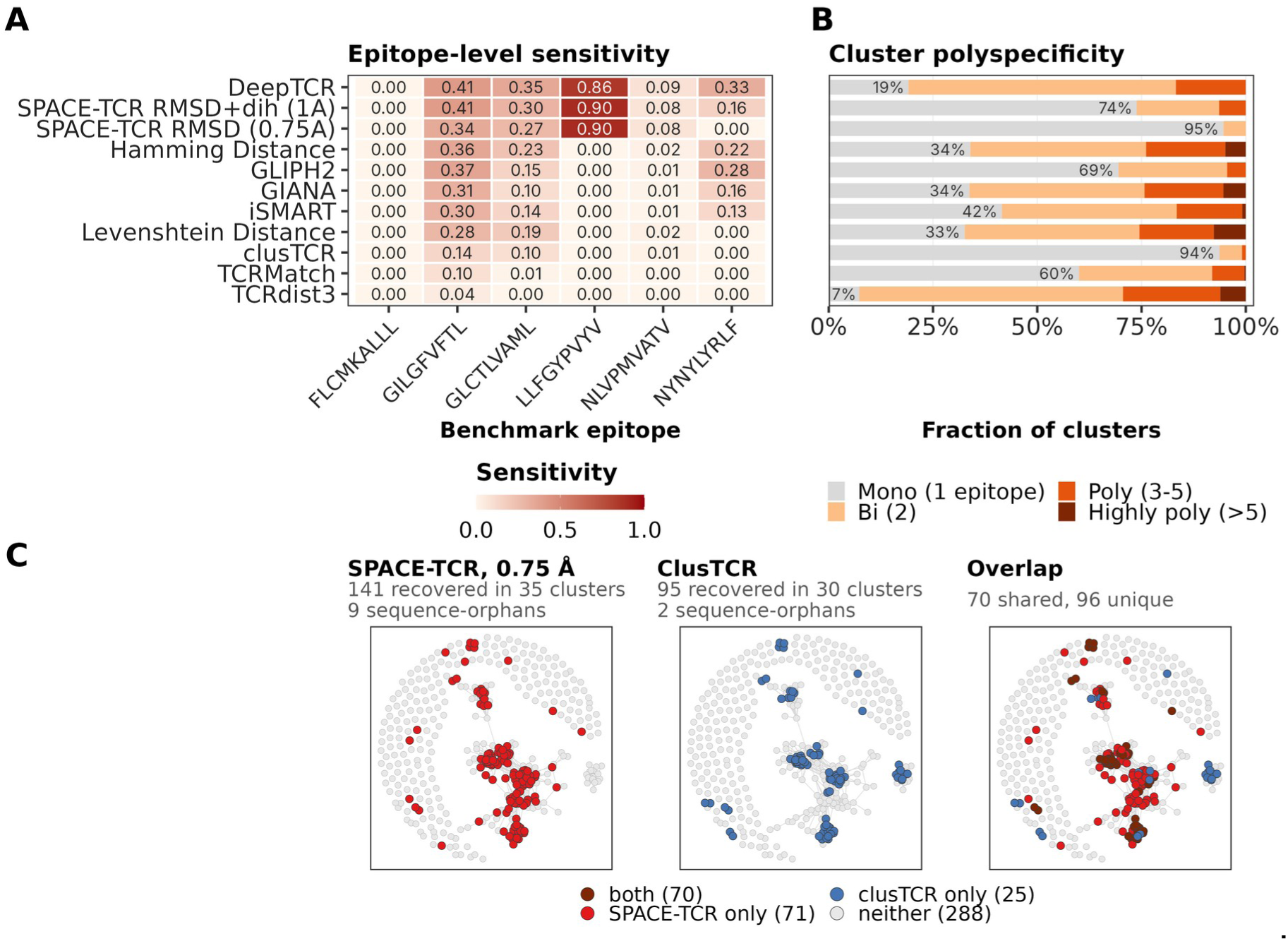
Epitope-level sensitivity and polyspecificity. (A) Sensitivity for eleven settings (nine published methods and two SPACE-TCR modes) across six benchmark epitopes using the Jouannet qualifying rule (cluster size > 3 and dominant fraction ≥ 67%). Grey cells denote zero recovered sensitivity, including sparsely represented epitopes. (B) Fractions of mono-, bi-, poly-, and highly polyspecific clusters. More than 94% of focal SPACE-TCR clusters are monospecific. (C) Coverage of 454 unique GILGFVFTL CDR3β sequences on a shared sequence graph, with an edge joining two sequences at Levenshtein distance ≤ 2 and drawn as three panels on one shared layout (SPACE-TCR, ClusTCR and their overlap), with each node coloured by which methods recovered it: 141 by SPACE-TCR and 95 by ClusTCR, 70 shared and 288 by neither. The receptors SPACE-TCR recovers alone lie largely in sparse and disconnected regions of the graph, whereas those ClusTCR recovers alone are concentrated in one dense neighbourhood.

Across the full cohort, sensitivity was strongly constrained by the number of receptors available for each epitope. The 367 epitopes hold a median of three unique CDR3β each: only 148 (40.3%) reach the four receptors a qualifying cluster requires, 47 (12.8%) reach ten, and 15 (4.1%) reach fifty. Sensitivity is therefore undefined in practice for most of the cohort, and every method recovers only a small minority of it. A qualifying cluster is formed for 22 of the 367 epitopes by DeepTCR, 18 by Hamming, 16 by GIANA, 15 by Levenshtein, 14 by iSMART and by SPACE-TCR under the gate, 12 by SPACE-TCR at the focal threshold, 10 by GLIPH2, seven by TCRMatch, four by ClusTCR and two by tcrdist3. Where an epitope is recovered at all, the median sensitivity is 0.291 for DeepTCR, 0.258 for SPACE-TCR under the gate, 0.206 for Hamming and 0.201 for SPACE-TCR at the focal threshold. Averaged over all 367 epitopes every method falls below 0.02.

SPACE-TCR averaged over the six epitopes ranks second and third of the eleven settings, behind only DeepTCR, while retaining the lowest polyspecificity of all methods (5.4%, against 31% for GLIPH2 and 81% for DeepTCR), and the clusters it forms include the sequence-divergent families described in Section 3.4 that none of the CDR3β-similarity methods produces. Each method is scored over the input it was given, so what is compared is the fraction of its own labelled receptors that each recovers in qualifying clusters, not an absolute count.

Figure 3C compares coverage of 454 unique GILGFVFTL CDR3β sequences on a common sequence graph, with edges joining receptors at Levenshtein distance ≤ 2. ClusTCR clustered 95 receptors (20.9%), principally in dense sequence neighbourhoods; SPACE-TCR clustered 141 (31.1%), including peripheral and disconnected receptors. Seventy were recovered by both methods, 71 only by SPACE-TCR, 25 only by ClusTCR, and 288 by neither. Structural clustering therefore reached beyond the sequence-dense cores rather than merely enlarging them.

### 3.6 Within-cluster sequence diversity and monospecificity

The published sequence methods were concentrated at median Levenshtein distance 0-1, as expected from their clustering criteria. SPACE-TCR at 0.75 Å had a median of 2 and a right tail extending to 8 (Figure 4). This distribution is the quantitative signature of the sequence-diverse and sequence-unrelated classes in Table 3. Two paired methods also form such clusters, and the comparison is therefore made on purity rather than count: DeepTCR produced 152 sequence-divergent high-purity clusters and tcrdist3 35, against 32 for SPACE-TCR at the focal threshold, but only 19% of DeepTCR’s 1,628 clusters and 7% of tcrdist3’s 669 reached purity ≥ 0.9, compared with 95% of SPACE-TCR’s 149 (global purity 0.63 and 0.42 against 0.99). SPACE-TCR is thus the only method that recovers sequence-divergent families without a corresponding loss of purity, and no CDR3β-similarity method recovers them at all. Apart from DeepTCR, SPACE-TCR was clearly separated from the sequence methods, consistent with cluster membership being driven by structural rather than motif similarity.

**Figure 4.**
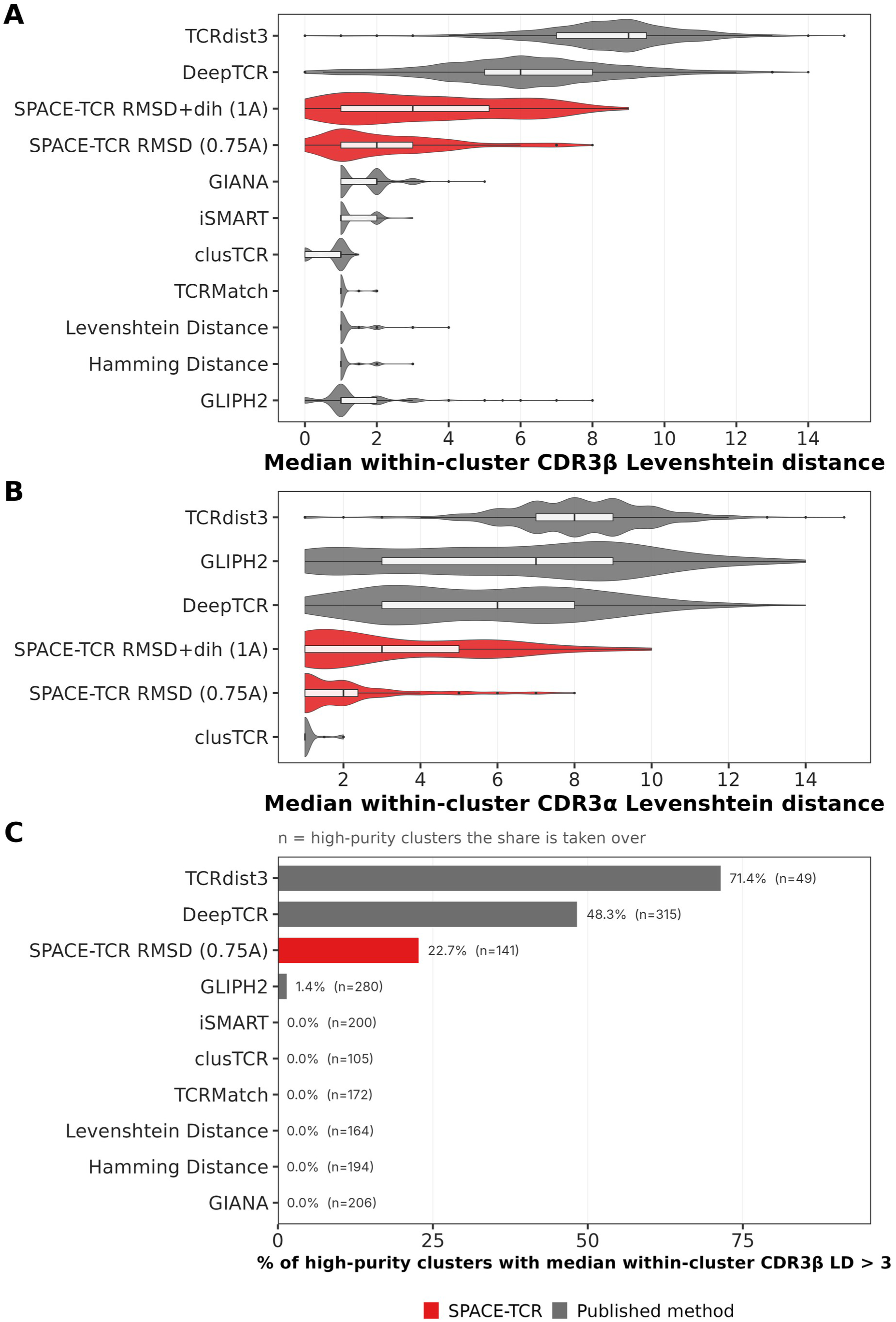
Within-cluster CDR3 sequence diversity. Violin plots show the distribution of the median pairwise Levenshtein distance per cluster. (A) CDR3β: sequence methods are concentrated near LD 0-1, whereas focal SPACE-TCR has median 2 and extends to 8. (B) CDR3α among methods with paired-chain output: SPACE-TCR and ClusTCR retain low within-cluster CDR3α sequence variation, whereas DeepTCR, GLIPH2, and tcrdist3 show greater variation. (C) Share of each method’s high-purity clusters (purity ≥ 0.9, size ≥ 2) whose median within-cluster CDR3β Levenshtein distance exceeds 3 — groups no edit-distance criterion would have formed. n is the number of high-purity clusters the share is taken over and should be read beside it, since a high percentage over few clusters is not a strong result.

The CDR3α comparison is informative because every paired-output method was supplied with both chains (Figure 4B). SPACE-TCR clusters retained low within-cluster CDR3α sequence variation: median within-cluster CDR3α Levenshtein distance was 2 (IQR 1-2.4; 114 clusters) at 0.75 Å and 3 (IQR 1-5; 145 clusters) for the 1.0 Å dihedral-gated mode. The distributions were much broader for tcrdist3 (median 8, IQR 7-9; 653 clusters), GLIPH2 (7, IQR 3-9; 468 clusters), and DeepTCR (6, IQR 3-8; 1,587 clusters). These methods produced greater within-cluster α-chain sequence variation despite receiving paired-chain input.

ClusTCR was the exception, with median CDR3α LD of 1, an interquartile range of 0, and a maximum of 2 among the 54 clusters containing any α-chain variation. This coherence is consistent with the stringency of its β-chain sequence criterion: receptors that are near-clonal in β are often near-clonal in α. SPACE-TCR achieved similar α-chain coherence across clusters whose CDR3β sequences spanned LD 0-8.

As SPACE-TCR clusters the paired αβ receptor, it can distinguish paired families within a single epitope. For NLVPMVATV, the focal mode resolved four recurrent αβ motif combinations (Figure S5): CAVRD… ARLMF with CASSS…NEAFF; CAGPMK…TSYDKVIF with CASSSA…YGYTF; CARNTGNQFYF with CASS… TGT…YGYTF; and CILDNNNDMRF with CASSLAPG…TNEKLFF. The α and β motifs co-varied within clusters, indicating distinct paired clonotype families rather than a shared β chain with interchangeable α partners. A CDR3β-only method cannot represent this joint structure and TCRBuilder2+ predicts the αβ Fv jointly. Replacing each receptor’s α chain with the same fixed α-chain sequence changed the predicted CDR3β backbone dihedral angles by a median of 0.692 rad, despite leaving the β-chain sequence unchanged. This change was 83% of the median difference between the β loops of two different receptors (0.835 rad; Figure S9D). When all receptors were modelled with the same α chain and clustering used only β-chain structure, purity decreased from 0.964 to 0.432 (Figure S9).

Polyspecificity provides a second measure of cluster contamination. At the focal threshold, 5.4% of SPACE-TCR clusters contained more than one epitope, the lowest proportion among the evaluated methods. ClusTCR was next at 6.3%; GLIPH2 and TCRMatch were 30.5% and 40.0%; iSMART, Hamming, GIANA, and Levenshtein ranged from 58.5% to 67.4%; DeepTCR was 80.9%; and tcrdist3 was 92.7% (Figure 3B). Structural clustering therefore combined high purity with low residual polyspecificity.

### 3.7 Germline gene composition: V-gene homogeneity without J-gene constraint

V-gene and J-gene homogeneity measures the clustering tools reliance on germline signals in their clustering behaviour (Figure 5). For V-gene homogeneity, SPACE-TCR at 0.75 Å and ClusTCR were the leading methods: 90% and 92% of clusters, respectively, had one V gene accounting for at least 90% of members. TCRMatch reached 69%, GLIPH2 62%, the distance baselines, GIANA, and DeepTCR 42-55%, and tcrdist3 5%. SPACE-TCR received no gene annotation, yet its clusters contained a mean of only 1.13 distinct V genes. This suggests that CDR3 loops superimposing within 0.75 Å are usually presented from the same germline framework.

**Figure 5.**
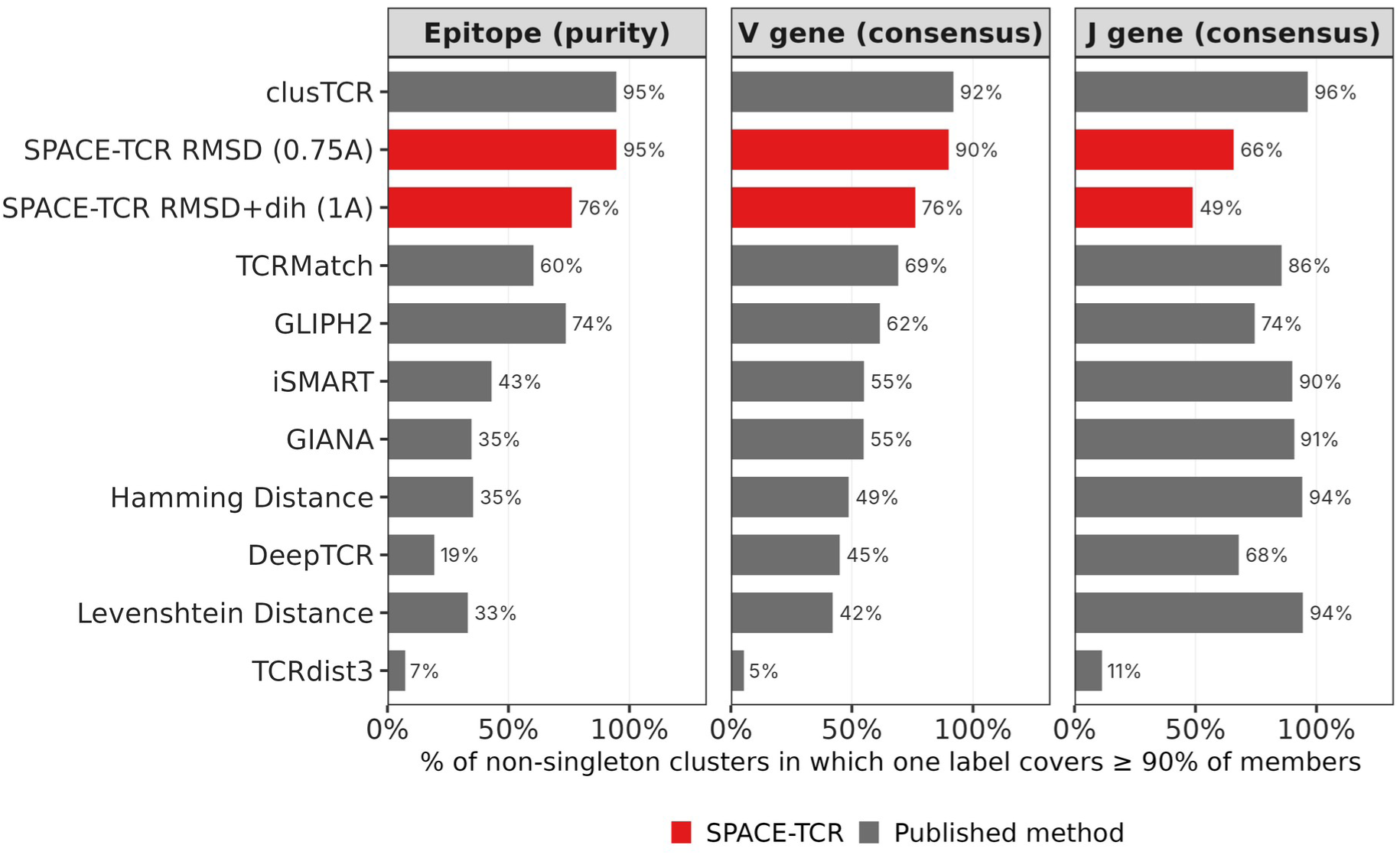
Epitope purity and TRBV/TRBJ homogeneity. Bars show the percentage of non-singleton clusters in which one label accounts for at least 90% of members. SPACE-TCR and ClusTCR combine high epitope purity with high V-gene homogeneity. Sequence-distance baselines show high J-gene homogeneity but much lower epitope purity. Gene homogeneity for pooled single-chain methods is calculated only in clusters containing at least two CDR3β sequences; epitope values use one label per receptor and therefore differ from the expanded multi-label values in Table 2.

However, the J-gene pattern was reversed. Hamming and Levenshtein reached 94% J-gene homogeneity, GIANA 91%, iSMART 90%, and TCRMatch 86%, whereas SPACE-TCR reached 66% at 0.75 Å and 49% in the 1.0 Å dihedral-gated mode. This pattern is expected because TRBJ contributes conserved C-terminal residues to CDR3β, so string distance partly reflects J-gene identity. Sequence-based methods can therefore be highly J-homogeneous without being epitope-pure. SPACE-TCR showed the opposite profile: high epitope and V-gene coherence with greater tolerance of J-gene variation. ClusTCR was high on all three measures, consistent with its near-clonal clusters.

### 3.8 Training-data leakage and held-out validation

We next tested whether TCRBuilder2+ training-set overlap could inflate performance. STCRDab (Leem et al., 2018), the structural T-cell receptor database from which TCRBuilder2+’s training set is drawn, contains 608 PDB entries with 279 distinct human CDR3β sequences. Forty-five of these sequences occurred in the benchmark: 45 of the cohort’s 4,292 unique CDR3β sequences (1.05%). Per-epitope overlap is reported over each epitope’s own sequence count and is therefore governed by the rare epitopes: four epitopes represented by one or three sequences lie entirely within STCRDab, whereas among the 47 epitopes with at least ten unique CDR3β the maximum is 10.0% and the median 0%. For the six benchmark epitopes the overlap is 1.8% or below, except the sparsely represented LLFGYPVYV at 1 of 7 (Figure S6A). Removing every modelled TCR carrying an overlapping CDR3β left 4,648 of the original 4,779 TCRs. At 0.75 Å, the held-out subset had 12.6% retention, 98.5% global purity, and 94.1% of clusters at purity ≥ 0.9, compared with 14.2%, 98.7%, and 94.6% in the full cohort (Figure S6B). The small change is consistent with the removal of 131 inputs and does not indicate a material memorisation effect.

### 3.9 Robustness to repertoire noise

The preceding analyses used a fully annotated cohort. To test performance when antigen-labelled receptors are a minority, we applied the series of noise levels described in Section 2.6 (Figure 6). On the undiluted subset of 10X dextramer labelled TCRs, the leading methods were close: SPACE-TCR reached 96.7% purity at the focal 0.75 Å threshold, compared with 94.7% for iSMART, 94.6% for GIANA, 94.0% for TCRMatch, 92.5% for GLIPH2, and 87.5% for ClusTCR.

**Figure 6.**
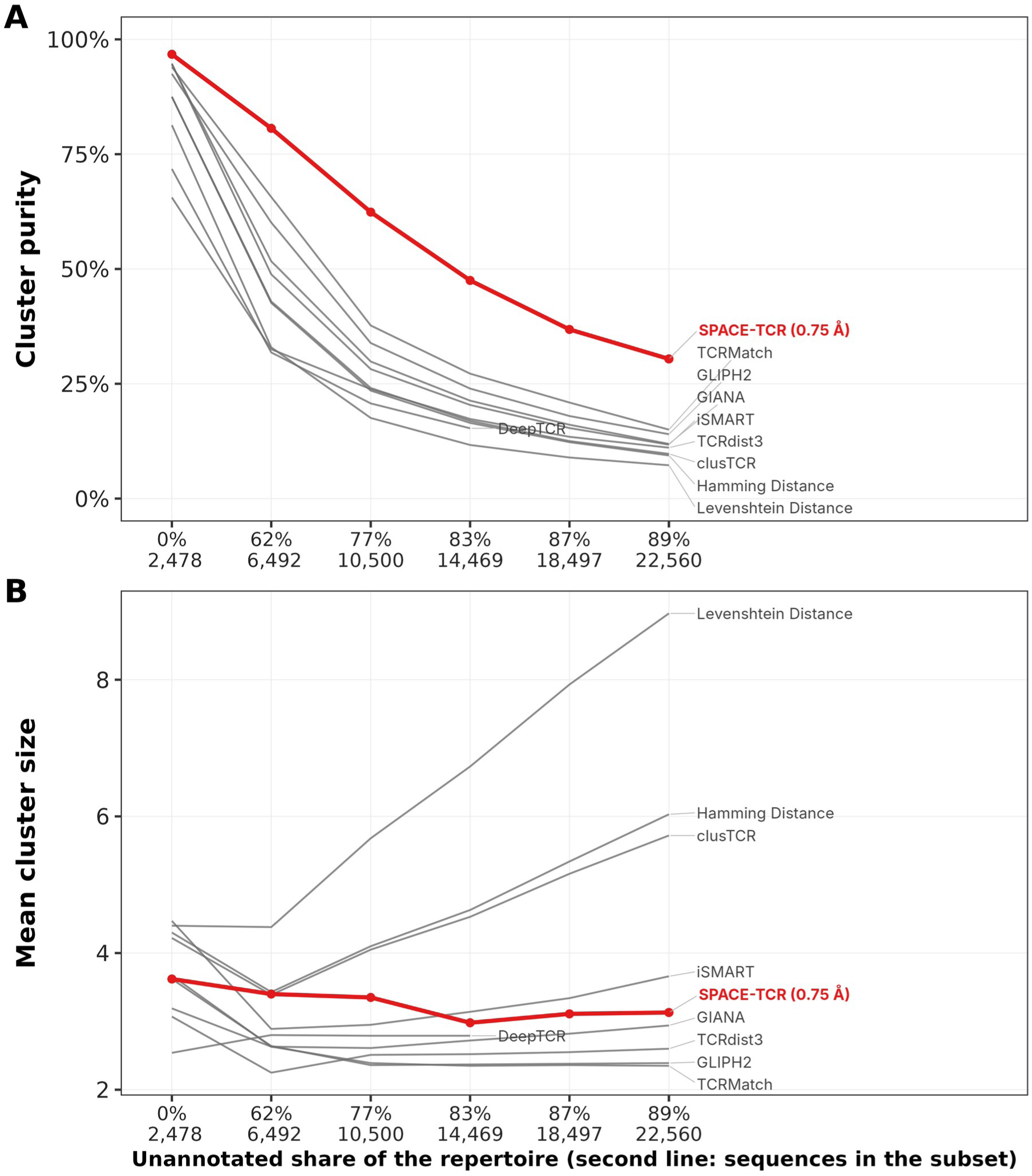
Robustness to repertoire noise across six nested 10X donor-1 subsets. Purity declines as unannotated members dilute cluster labels; the relevant comparison is the rate of decline. SPACE-TCR at 0.75 Å has the highest purity at every noise level. The margin over the best published method widens as noise rises: 96.7% against iSMART’s 94.7% at the lowest noise level, and 30.4% against TCRMatch’s 15.0% at the highest noise level.

As unannotated ‘noise’ TCRs were progressively introduced, the methods separated clearly. At the focal 0.75 Å threshold, SPACE-TCR had the highest purity at every noise level and maintained 30.4% purity at 89.2% unannotated content. The best published comparator at that point was TCRMatch at 15.0%, followed by GLIPH2 at 14.0%, GIANA at 11.9%, iSMART at 11.7%, ClusTCR at 9.7%, and the Levenshtein baseline at 7.3%. The tighter 0.5 Å SPACE-TCR setting maintained 62.4% purity. Relative to their zero-noise values, the focal and 0.5 Å settings retained 31% and 65% of their initial purity, respectively, whereas the published methods retained 8-16%. Thus, purity on the undiluted subset did not predict robustness to repertoire noise.

Changes in mean cluster size help to explain the decline. From zero to 89.2% noise, mean size increased from 4.4 to 9.0 for the Levenshtein baseline, from 4.3 to 6.0 for Hamming, and from 4.2 to 5.7 for ClusTCR. These methods use absolute edit-distance thresholds, so additional sequences can connect to and enlarge clusters that already contain annotated signal. In contrast, SPACE-TCR mean size decreased from 3.6 to 3.0 at 0.5 Å and from 3.6 to 3.1 at 0.75 Å. The added sequences therefore formed separate small clusters more often than they contaminated existing ones. GLIPH2 and GIANA also decreased in mean size, indicating that this absorption pattern is specific to the absolute sequence-distance methods rather than to all sequence-based approaches.

### 3.10 Consensus clustering with sequence-based methods

As structural and sequence methods recovered partly different cluster populations, we tested whether agreement between them improved cluster quality. Consensus clusters required both SPACE-TCR and the sequence method to group a receptor pair (Section 2.5). On the shared universe of 6,163 unique CDR3β sequences, all four consensus approaches exceeded both parents in the proportion of clusters at purity ≥ 0.9 (Figure S7A). The two GLIPH2 consensuses reached 99%, compared with 73% for GLIPH2 alone on this universe (51.0% on the 4,292-sequence set of Table 2); the two ClusTCR consensuses reached 98%, compared with 96% for ClusTCR. The largest improvements occurred when one parent had lower purity, consistent with the two representations making partly independent errors.

The improvement came at the expected cost in yield (Figure S7B). The four consensuses reached 99.4-99.7% global purity over 2.2-4.4% retention, compared with 98.7% purity at 7.2% retention for SPACE-TCR at 0.75 Å, 98.8% at 3.3% for ClusTCR, and 90.2% at 11.6% for GLIPH2. Consensus clustering is therefore suited to settings in which each candidate is expensive to validate and a small, highly confident set is preferred. It is less suitable for discovery because intersection with a sequence method necessarily removes the sequence-diverse clusters that motivate the structural analysis.

### 3.11 Re-validation against functionally screened epitope labels

Following analysis on the Jouannet benchmark, we applied SPACE-TCR to the functionally screened TCR data released from TCRvdb on 5 epitopes from their study (Messemaker et al., 2026). Of 1,514 TCRs, 1,378 matched a TCRvdb entry, and among the 1,151 with an unambiguous call only 557 (48.4%) validated. Cluster purity nevertheless proved robust: at the focal 0.75 Å operating point it fell from 99.6% to 95.7% when every non-validating receptor was relabelled to a class of its own. Over the same 1,151 receptors SPACE-TCR clustered twice as many as ClusTCR (20.7% versus 10.3%) at comparable purity. Moreover, receptors that SPACE-TCR clustered validated experimentally considerably more often than those it left unclustered (79% versus 28% among receptors with no close sequence relatives defined by 0 sequence neighbours with CDR3 LD <2), while ClusTCR showed no significant difference on the same receptors (Supplementary Section S19, Figure S8).

## 4. Discussion

This benchmark supports three principal conclusions. First, SPACE-TCR performs at the high-purity end of the published methods despite clustering on predicted three-dimensional CDR3 geometry rather than sequence. At the focal threshold, 94.6% of clusters had purity ≥ 0.9, second only to ClusTCR (97.3%), while SPACE-TCR clustered substantially more TCRs. It also had the lowest polyspecificity (5.4%). These results indicate that paired structural information is at least comparable with sequence for forming high-confidence specificity groups.

Secondly, the structural representation formed high-purity clusters that CDR3β-similarity criteria would not form: 22.7% of high-purity focal clusters were sequence-diverse or sequence-unrelated, increasing to 40% at 1.0 Å. As the method operates on the paired receptor, it also separated single epitopes into coherent αβ families in which the two chain motifs co-varied. Thirdly, excluding all TCRBuilder2+ training-set-overlapping CDR3β sequences changed the principal purity measures by less than one percentage point, arguing against training-set memorisation as the explanation for the benchmark performance.

Huang et al. (2026) predicted paired TCR structures with AlphaFold3 and tFold-TCR, clustered them with Foldseek and with CDR3 structural embeddings, and reported more antigen-specific clusters than sequence clustering recovered, with two discovered receptors validated experimentally; their evaluation covers ten peptides and one sequence comparator. Mayol-Rullan et al. (2025) encoded modelled CDR loops as physicochemical fingerprints and clustered TCR repertoires from them, but found their structural representation underperformed a sequence baseline (29.8% against 41.4% rank-1 same-specificity pairing), reaching high accuracy only above a similarity cut-off that retained 12.3% of receptors. The present work differs from both in representation—Cα RMSD across the CDR loops after framework superposition—and in scope: 367 epitopes against ten, nine published comparators against one, and the purity, retention and sensitivity definitions of an existing benchmark.

SPACE-TCR clusters were highly homogeneous in V-gene usage, although germline annotations were not supplied as clustering inputs. They were less homogeneous in J-gene usage than clusters formed by edit-distance methods. This pattern is consistent with structural similarity capturing the presentation of the antigen-contacting loop rather than simply the J-encoded CDR3β stem. It is also consistent with where predicted TCR structures are reliable: CDR3 confidence correlates with CDR3 accuracy more strongly than for any other region, whereas the orientation of the receptor over the peptide–MHC is the dominant source of error (Lu et al., 2026). SPACE-TCR clusters the unbound receptor’s CDR loops after framework alignment and therefore does not depend on the docking geometry that limits complex prediction. Consensus clustering further demonstrated complementarity: intersecting SPACE-TCR with ClusTCR or GLIPH2 increased high-purity cluster rates to 98-99%, but reduced retention and removed the sequence-diverse structural clusters.

The noise benchmark is particularly relevant to clinical repertoires. At the focal 0.75 Å threshold, SPACE-TCR had the highest purity at every noise level and remained at 30.4% purity with 89.2% unannotated content, compared with 15.0% for the best published comparator. The 0.5 Å setting reached 62.4%. The associated cluster-size trends suggest that absolute edit-distance methods progressively absorb unannotated neighbours into signal-bearing clusters, whereas SPACE-TCR tends to keep added sequences in separate small clusters.

These findings suggest several applications for structural clustering in repertoire analysis. By recovering sequence-divergent receptors with potentially shared specificity, SPACE-TCR could facilitate the discovery of antigen-reactive TCRs and expand the pool of candidates for therapeutic development. More broadly, structural receptor families could provide a biologically informative unit for comparing immune responses across individuals, identifying disease-associated repertoires, and monitoring changes following infection, vaccination or treatment. Such comparisons are complicated by the fact that repertoire composition itself varies with HLA genotype, which shapes both the receptors recruited to an epitope and their cross-reactive memory potential (Francis et al., 2022); a unit defined on receptor geometry rather than on a shared sequence motif does not presuppose that the same motif recurs across individuals. The ability to recover high-purity clusters from largely unannotated backgrounds is particularly relevant to these applications, where antigen labels are generally unavailable. Although these applications require validation in independent clinical cohorts, the present findings suggest that structural convergence could reveal aspects of antigen-driven repertoire organisation that conventional sequence-based approaches fail to capture.

Several limitations constrain these conclusions. The focal range of 0.75-1.0 Å is specific to the SPACE-TCR backbone representation and TCRBuilder2+ model quality, and would require recalibration for another predictor such as TCRmodel2 (Yin et al., 2023). It is also smaller than the absolute accuracy of current CDR3 predictions. For the TCR-specific predictor most comparable to the one used here, TCRBuilder2+, median backbone errors are below 2 Å across every CDR loop, and the two antigen-contacting loops differ: 2 of 45 held-out receptors fall outside 2 Å for CDR3β against 13 of 45 for CDR3α (Quast et al., 2025). The best reported antibody-domain predictors sit in the same 1.8-1.9 Å range for CDR3 (Dreyer et al., 2025). The two quantities are not directly comparable: clustering compares models with each other after framework alignment rather than against experimental structures, so systematic error shared across models of similar receptors largely cancels, and only the relative geometry has to be reliable. The empirical results are consistent with this — 98.7% purity at the focal threshold, and an unchanged result when training-set-overlapping receptors are removed — but it does mean the thresholds reported here should be read as properties of this model ensemble rather than as physical distances. Quast et al. (2025) further report that only about one third of modelled TCRs are predicted accurately enough for reliable analysis of the receptor-peptide interface. SPACE-TCR does not use the interface: it compares the relative geometry of the unbound CDR loops after framework alignment, which is the less demanding requirement of the two.

Structure-guided analysis is also substantially more computationally expensive than direct sequence clustering. TCRBuilder2+ predicts a paired structure in under one minute on CPU or about five seconds on GPU, compared with roughly 30 minutes for AlphaFold-Multimer (Abanades et al., 2023), but modelling millions of receptors remains the dominant throughput constraint, although recent predictors have improved both prediction speed and CDR-loop accuracy. Dedicated immune-protein predictors continue to get faster — ABodyBuilder3 reports a threefold speed-up over its predecessor (Kenlay et al., 2024), and the pan-immunoglobulin model Ibex predicts a variable domain in about 0.7 s on a single GPU, roughly two orders of magnitude faster than general-purpose structure predictors (Dreyer et al., 2025) — and TCR-specific models have improved on precisely the regions this method uses: TCRBuilder2+, retrained on an expanded set of solved TCR structures, reports median errors below 2 Å across the CDR loops of both chains, with CDR3β the better resolved (Quast et al., 2025). Predicted structures are also now produced at scale by others: Quast et al. (2025) released approximately 1.5 million predicted TCR structures, and a further 14,612 paired TCR models accompanied the 2026 VDJdb release (Luppov et al., 2026). Faster prediction and more accurate CDR-loop models should make structure-based repertoire analysis more practical.

We tested whether the clustering could be distilled into a sequence-only model, so that structures would be needed only for candidate groups: a contrastive encoder trained on the SPACE-TCR clusters did not reproduce them at the focal threshold and did not transfer to an independent repertoire, so the constraint stands (Supplementary Note S12). The encoder was trained from scratch on the cohort rather than initialised from a pretrained immune-receptor language model. Domain-specific pretraining of this kind is well established for antibodies, where AbLang (Olsen et al., 2022) and its paired-chain successor AbLang2 learn representations from large repertoire corpora. An analogous pretraining approach could be evaluated using a larger paired TCR dataset. Structural modelling of the TCR-pMHC complex cannot yet be used to infer specificity reliably from a predicted interface. The question this work asks is what can be recovered without it. Modelling the unbound receptor alone, a task current predictors perform comparatively well, and comparing the relative geometry of the CDR loops is sufficient to form groups that are as pure as the best sequence methods and that sequence similarity would not form at all. Those groups are hypotheses about shared recognition rather than demonstrations of it. Testing them is becoming practical: deep peptide-recognition profiling now assigns specificities to receptors at scale and has been used to recover disease-associated antigens (Wang et al., 2026), and a structural grouping offers a principled way to choose which receptors to profile. These results show that paired structural models can reveal specificity-associated receptor groups that are not recovered by the sequence-based criteria evaluated here.

### Summary of key findings

1. **High cluster purity.** At the focal threshold, 94.6% of SPACE-TCR clusters had purity ≥ 0.9, compared with 97.3% for ClusTCR and no more than 63.5% for any other method.
2. **Greater yield than the only comparably pure method.** SPACE-TCR clustered 446 of the cohort’s 4,292 unique CDR3β (10.4%) in 149 clusters, against ClusTCR’s 201 (4.7%) in 111.
3. **Structural clusters beyond standard sequence criteria.** At 0.75 Å, 22.7% of high-purity clusters were sequence-diverse or sequence-unrelated; the proportion increased to 40.0% at 1.0 Å.
4. **Lowest polyspecificity.** Only 5.4% of focal SPACE-TCR clusters contained more than one epitope, compared with 6.3% for ClusTCR and more than 30% for every other method.
5. **Greater within-cluster CDR3β diversity.** The median within-cluster Levenshtein distance was 2, with values extending from 0 to 8, compared with a median of 1 for the sequence methods.
6. **Robustness to training-set overlap.** Removing every modelled TCR whose CDR3β sequence occurred in STCRDab changed the principal purity metrics by less than one percentage point.
7. **V-gene homogeneity without germline input.** One V gene accounted for at least 90% of members in 90% of focal clusters, compared with 92% for ClusTCR, although SPACE-TCR received no gene annotation.
8. **Paired-chain coherence across sequence-divergent receptors.** Median within-cluster CDR3α Levenshtein distance was 2 for SPACE-TCR, compared with 6 for DeepTCR and 8 for tcrdist3 despite paired input.
9. **Higher confidence through consensus.** Intersecting SPACE-TCR with ClusTCR or GLIPH2 produced 98-99% high-purity clusters, at the expected cost in retention.
10. **Highest purity under repertoire noise at the focal threshold.** SPACE-TCR at 0.75 Å had higher purity than the evaluated sequence-based methods across all available comparisons, maintaining 30.4% at 89.2% unannotated content against 15.0% for the best published comparator; the tighter 0.5 Å mode, which clusters far fewer receptors, maintained 62.4%.

## References

Abanades, B., Wong, W. K., Boyles, F., Georges, G., Bujotzek, A., & Deane, C. M. (2023). ImmuneBuilder: Deep-Learning models for predicting the structures of immune proteins. Communications Biology, 6(1), 575. 10.1038/s42003-023-04927-7

Chronister, W. D., Crinklaw, A., Mahajan, S., Vita, R., Koşaloğlu-Yalçn, Z., Yan, Z., Greenbaum, J. A., Jessen, L. E., Nielsen, M., Christley, S., Cowell, L. G., Sette, A., & Peters, B. (2021). TCRMatch: predicting T-cell receptor specificity based on sequence similarity to previously characterized receptors. Frontiers in Immunology, 12, 640725. 10.3389/fimmu.2021.640725

Dash, P., Fiore-Gartland, A. J., Hertz, T., Wang, G. C., Sharma, S., Souquette, A., Crawford, J. C., Clemens, E. B., Nguyen, T. H. O., Kedzierska, K., La Gruta, N. L., Bradley, P., & Thomas, P. G. (2017). Quantifiable predictive features define epitope-specific T cell receptor repertoires. Nature, 547(7661), 89–93. 10.1038/nature22383

Dreyer, F. A., Ludwiczak, J., Martinkus, K., Abanades, B., Alberstein, R. G., Kessel, P., Rao, P., Lee, J. H., Bonneau, R., Watkins, A. M., & Seeger, F. (2025). Conformation-aware structure prediction of antigen-recognizing immune proteins. mAbs, 18(1), 2602217. 10.1080/19420862.2025.2602217

Francis, J. M., Leistritz-Edwards, D., Dunn, A., Tarr, C., Lehman, J., Dempsey, C., Hamel, A., Rayon, V., Liu, G., Wang, Y., Wille, M., Durkin, M., Hadley, K., Sheena, A., Roscoe, B., Ng, M., Rockwell, G., Manto, M., Gienger, E., … Pregibon, D. C. (2022). Allelic variation in class I HLA determines CD8+ T cell repertoire shape and cross-reactive memory responses to SARS-CoV-2. Science Immunology, 7(67), eabk3070. 10.1126/sciimmunol.abk3070

Glanville, J., Huang, H., Nau, A., Hatton, O., Wagar, L. E., Rubelt, F., Ji, X., Han, A., Krams, S. M., Pettus, C., Haas, N., Lindestam Arlehamn, C. S., Sette, A., Boyd, S. D., Scriba, T. J., Martinez, O. M., & Davis, M. M. (2017). Identifying specificity groups in the T cell receptor repertoire. Nature, 547(7661), 94–98. 10.1038/nature22976

Hamming, R. W. (1950). Error detecting and error correcting codes. Bell System Technical Journal, 29(2), 147–160. 10.1002/j.1538-7305.1950.tb00463.x

Heather, J. M., Spindler, M. J., Herrero Alonso, M., Shui, Y. I., Millar, D. G., Johnson, D. S., Cobbold, M., & Hata, A. N. (2022). Stitchr: stitching coding TCR nucleotide sequences from V/J/CDR3 information. Nucleic Acids Research, 50(12), e68. 10.1093/nar/gkac190

Huang, H., Wang, C., Rubelt, F., Scriba, T. J., & Davis, M. M. (2020). Analyzing the Mycobacterium tuberculosis immune response by T-cell receptor clustering with GLIPH2 and genome-wide antigen screening. Nature Biotechnology, 38(10), 1194–1202. 10.1038/s41587-020-0505-4

Huang, Q., Liu, Y., Tang, X., Zhang, W., Cao, Y., Zhao, J., Qi, F., & Zhang, Z. (2026). A comparative and exploratory analysis of computational methods for TCR structural prediction and antigen-specific TCR discovery. Briefings in Bioinformatics, 27(4), bbag382. 10.1093/bib/bbag382

Hudson, D., Lubbock, A., Basham, M., & Koohy, H. (2024). A comparison of clustering models for inference of T cell receptor antigen specificity. ImmunoInformatics, 13, 100033. 10.1016/j.immuno.2024.100033

Jouannet, C., Vantomme, H., Le Gouge, K., Klatzmann, D., & Mariotti-Ferrandiz, E. (2025). Benchmarking unsupervised methods for inferring TCR specificity. NAR Genomics and Bioinformatics, 7(4), lqaf150. 10.1093/nargab/lqaf150

Kenlay, H., Dreyer, F. A., Cutting, D., Nissley, D., & Deane, C. M. (2024). ABodyBuilder3: improved and scalable antibody structure predictions. Bioinformatics, 40(10), btae576. 10.1093/bioinformatics/btae576

Leem, J., de Oliveira, S. H. P., Krawczyk, K., & Deane, C. M. (2018). STCRDab: the structural T-cell receptor database. Nucleic Acids Research, 46(D1), D406–D412. 10.1093/nar/gkx971

Levenshtein, V. I. (1966). Binary codes capable of correcting deletions, insertions, and reversals. Soviet Physics Doklady, 10(8), 707–710.

Lu, J., Zhu, X., Hu, X., Zhang, C., & Feng, F. (2026). Benchmarking TCR–pMHC structure prediction: a unified evaluation and CDR3-based functional insights. Briefings in Bioinformatics, 27(3), bbag289. 10.1093/bib/bbag289

Luppov, D. V, Koneva, A. E., Bagaev, D. V, Alexandrova, A. V, Vlasova, E. K., Chudakov, D. M., Motozono, C., Sewell, A. K., & Shugay, M. (2026). VDJdb in 2026: boosting T-cell receptor recognition evidence using paratope embeddings and AI-based structure prediction. Nucleic Acids Research, gkag904. 10.1093/nar/gkag904

Mayer-Blackwell, K., Schattgen, S., Cohen-Lavi, L., Crawford, J. C., Souquette, A., Gaevert, J. A., Hertz, T., Thomas, P. G., Bradley, P., & Fiore-Gartland, A. (2021). TCR meta-clonotypes for biomarker discovery with tcrdist3: identification of public, HLA-restricted clusters of SARS-CoV-2 TCRs. eLife, 10, e68605. 10.7554/eLife.68605

Mayol-Rullan, F., Bugnon, L. A., Perez, M. A. S., & Zoete, V. (2025). A fingerprint approach to pioneer structure-based T cell receptor repertoire analysis and specificity prediction. Frontiers in Immunology, 16, 1688805. 10.3389/fimmu.2025.1688805

McMaster, B., Thorpe, C. J., Rossjohn, J., Deane, C. M., & Koohy, H. (2024). Quantifying conformational changes in the TCR:pMHC-I binding interface. Frontiers in Immunology, 15, 1491656. 10.3389/fimmu.2024.1491656

Messemaker, M., Kwee, B. P. Y., & Moravec, Ž. (2026). Functional evaluation of TCR-pMHC pairs at scale allows in silico TCR reactivity prediction. Immunity, 59, 2775–2788. 10.1016/j.immuni.2026.09.007

Nagano, Y., Pyo, A. G. T., Milighetti, M., Henderson, J., Shawe-Taylor, J., Chain, B., & Tiffeau-Mayer, A. (2025). Contrastive learning of T cell receptor representations. Cell Systems, 16(1), 101165. 10.1016/j.cels.2024.12.006

North, B., Lehmann, A., & Dunbrack, R. L. (2011). A new clustering of antibody CDR loop conformations. Journal of Molecular Biology, 406(2), 228–256. 10.1016/j.jmb.2010.10.030

Olsen, T. H., Moal, I. H., & Deane, C. M. (2022). AbLang: an antibody language model for completing antibody sequences. Bioinformatics Advances, 2(1), vbac046. 10.1093/bioadv/vbac046

Quast, N. P., Abanades, B., Guloglu, B., Karuppiah, V., Harper, S., Raybould, M. I. J., & Deane, C. M. (2025). T-cell receptor structures and predictive models reveal comparable alpha and beta chain structural diversity despite differing genetic complexity. Communications Biology, 8, 362. 10.1038/s42003-025-07708-6

Sidhom, J.-W., Larman, H. B., Pardoll, D. M., & Baras, A. S. (2021). DeepTCR is a deep learning framework for revealing sequence concepts within T-cell repertoires. Nature Communications, 12(1), 1605. 10.1038/s41467-021-21879-w

Spoendlin, F. C., Abanades, B., Raybould, M. I. J., Wong, W. K., Georges, G., & Deane, C. M. (2023). Improved computational epitope profiling using structural models identifies a broader diversity of antibodies that bind the same epitope. Frontiers in Molecular Biosciences, 10, 1237621. 10.3389/fmolb.2023.1237621

Tickotsky, N., Sagiv, T., Prilusky, J., Shifrut, E., & Friedman, N. (2017). McPAS-TCR: a manually curated catalogue of pathology-associated T cell receptor sequences. Bioinformatics, 33(18), 2924–2929. 10.1093/bioinformatics/btx286

Valkiers, S., Van Houcke, M., Laukens, K., & Meysman, P. (2021). ClusTCR: a Python interface for rapid clustering of large sets of CDR3 sequences with unknown antigen specificity. Bioinformatics, 37(24), 4865–4867. 10.1093/bioinformatics/btab446

Vita, R., Blazeska, N., Marrama, D., Shackelford, D., Zalman, L., Foos, G., Zarebski, L., Chan, K., Reardon, B., Fitzpatrick, S., Busse, M., Coleman, S., Sedwick, C., Edwards, L., MacFarlane, C., Ennis, M., Duesing, S., Bennett, J., Greenbaum, J., … Peters, B. (2025). The Immune Epitope Database (IEDB): 2024 update. Nucleic Acids Research, 53(D1), D436–D443. 10.1093/nar/gkae1092

Wang, N., Yeh, H., Lai, B., Perera, J., Jude, K. M., Risch, I., Um, J., Chen, X., Xiang, X., Wang, C., Liu, L. D., Yang, X., Paley, M. A., Khan, A. A., & Garcia, K. C. (2026). Deep peptide recognition profiling decodes TCR specificity and enables disease-associated antigen discovery. Nature Biotechnology. 10.1038/s41587-026-03128-x

Yin, R., Ribeiro-Filho, H. V, Lin, V., Gowthaman, R., Cheung, M., & Pierce, B. G. (2023). TCRmodel2: high-resolution modeling of T cell receptor recognition using deep learning. Nucleic Acids Research, 51(W1), W569–W576. 10.1093/nar/gkad356

Zhang, H., Liu, L., Zhang, J., Chen, J., Ye, J., Shukla, S., Qiao, J., Zhan, X., Chen, H., Wu, C. J., Fu, Y.-X., & Li, B. (2020). Investigation of antigen-specific T-cell receptor clusters in human cancers. Clinical Cancer Research, 26(6), 1359–1371. 10.1158/1078-0432.CCR-19-3249

Zhang, H., Zhan, X., & Li, B. (2021). GIANA allows computationally-efficient TCR clustering and multi-disease repertoire classification by isometric transformation. Nature Communications, 12(1), 4699. 10.1038/s41467-021-25006-7

