## Supplementary for "Structural clustering of paired αβ T-cell receptor repertoires recovers antigen-specific convergence beyond sequence similarity"

### S1. Cohort and clustering data trail

Table S1 traces the benchmark from the curated cohort through structural modelling to the reported clusters. Three denominators are used and should not be interchanged: 4,779 paired TCRs in the curated cohort (4,031 of them modelled by TCRBuilder2+), which underlie Tables 1 and 2 and the retention analyses in Sections 3.1-3.5; 4,292 unique CDR3β sequences within that cohort, used for the STCRDab-overlap analysis; and 6,163 unique CDR3β sequences in the shared consensus universe of Section 3.10, which spans all methods rather than the SPACE-TCR input alone. Thirteen CDR3β sequences in the cohort carry two epitope labels each, so per-epitope counts depend on how those are handled. Where per-epitope counts are required to sum to the 4,292 total, each shared sequence is assigned to one epitope, which gives 454 for GILGFVFTL and 345 for NLVPMVATV (Figures 2 and S6). Where an epitope is treated on its own, as in the sensitivity denominators of Section 3.5 and Supplementary Section S13, a shared sequence is counted under each of its labels, which gives 455 and 349. The two conventions differ by 13 sequences in total and by at most four for any one epitope.

**Supplementary Table S1.** Data trail from the curated cohort to the reported analyses.

| Stage | Count | Reported in |
|---|---|---|
| Paired TCRs from Jouannet | 4,779 | Table 1 |
| Unique CDR3β sequences within that input | 4,292 | Section 3.8 |
| Distinct epitopes represented in the curated cohort | 367 | Section 2.3 |
| Shared unique-CDR3β universe across all evaluated methods | 6,163 | Section 2.6 |
| Clusters formed at the focal 0.75 Å RMSD threshold | 149 | Table 1 |
| TCRs clustered at the focal threshold (rows; 14.2% retention) | 681 | Table 1 |
| Unique CDR3β among those clustered TCRs | 457 | Table S1b |
| Unique CDR3β in focal clusters of size ≥ 2 | 446 | Table S1b |
| High-purity clusters (purity ≥ 0.9, size ≥ 2) at the focal threshold | 141 | Section 3.4 |
| Distinct dominant epitopes across all focal-threshold clusters | 26 | Table S2 |
| Of those, also dominating a cluster at purity ≥ 0.9 | 26 | Table S2 |
| Of those, also dominating a high-purity cluster of ≥ 5 members | 9 | Table S2 |
| STCRDab reference set: PDB entries / unique human CDR3β strings | 608 / 279 | Section 3.8 |
| Unique input CDR3β sequences found in STCRDab | 45 (1.05%) | Section 3.8 |
| Modelled input TCRs carrying an STCRDab-matched CDR3β | 131 | Section 3.8 |
| Held-out input after STCRDab exclusion | 4,648 | Section 3.8 |

Table 2 reports the unique-CDR3β unit of Section 2.4 rather than clustered rows, because methods differ several-fold in the number of rows they emit per unique sequence and a row count is therefore not comparable between them. GLIPH2 produced 2,940 rows from 713 unique sequences (4.12 rows per sequence), ClusTCR 426 rows from 227 sequences (1.88), and SPACE-TCR 681 rows from 457 sequences (1.49). Restricting to clusters holding at least two sequences from the cohort's 4,292, as

Table 2 does, gives 446 for SPACE-TCR and 201 for ClusTCR. SPACE-TCR multiplicity arises when receptors share CDR3β but differ in CDR3α. On the unique-sequence unit, SPACE-TCR placed 446 sequences in non-singleton clusters compared with 201 for ClusTCR. The yield difference is therefore about twofold, rather than the 60% suggested by row counts.

Pooling all published methods, 6,162 unique CDR3β sequences were clustered by at least one published method; adding the one sequence clustered only by SPACE-TCR gives the 6,163-sequence shared universe. SPACE-TCR clustered 669 across its two focal modes, and 668 of these also appeared in a published method's output. Its advantage is therefore not unique receptor coverage. It lies in cluster composition: which receptors are grouped together, at what purity, and with what sequence diversity. The 22.7% sequence-unreachable estimate in Table 3 refers to cluster composition, not to receptors that no sequence method clusters at all.

**Supplementary Table S1b.** Clustered row counts and unique-CDR3β counts by method, showing the effect of overlapping cluster assignments.

| Method | Clustered rows | Unique CDR3β | Rows per unique CDR3β | Unique CDR3β in clusters ≥ 2 |
|---|---|---|---|---|
| GLIPH2 | 2,940 | 713 | 4.12 | 713 |
| SPACE-TCR RMSD+dih (1.0 Å) | 903 | 654 | 1.38 | 642 |
| SPACE-TCR RMSD (0.75 Å) | 681 | 457 | 1.49 | 446 |
| ClusTCR | 426 | 227 | 1.88 | 201 |

### S2. Distinct epitopes captured per method

Figure S1 and Table S2 assess epitope breadth at three quality thresholds: any non-singleton cluster, purity ≥ 0.9, and purity ≥ 0.9 with at least five members. DeepTCR reached 266 of the 367 epitopes, but only 48 remained after the purity filter and 14 after the size filter; tcrdist3 fell from 145 to 11 and then 2. SPACE-TCR at 0.75 Å reached 26 epitopes, all of which appeared in a high-purity cluster, and nine remained after the size filter. ClusTCR showed a similar profile (21, 19, and 5). The 1.0 Å dihedral-gated mode reached 40 epitopes, of which 25 were high-purity and 10 also met the size criterion. The strictest column is therefore the most relevant when nominating specificities for follow-up.

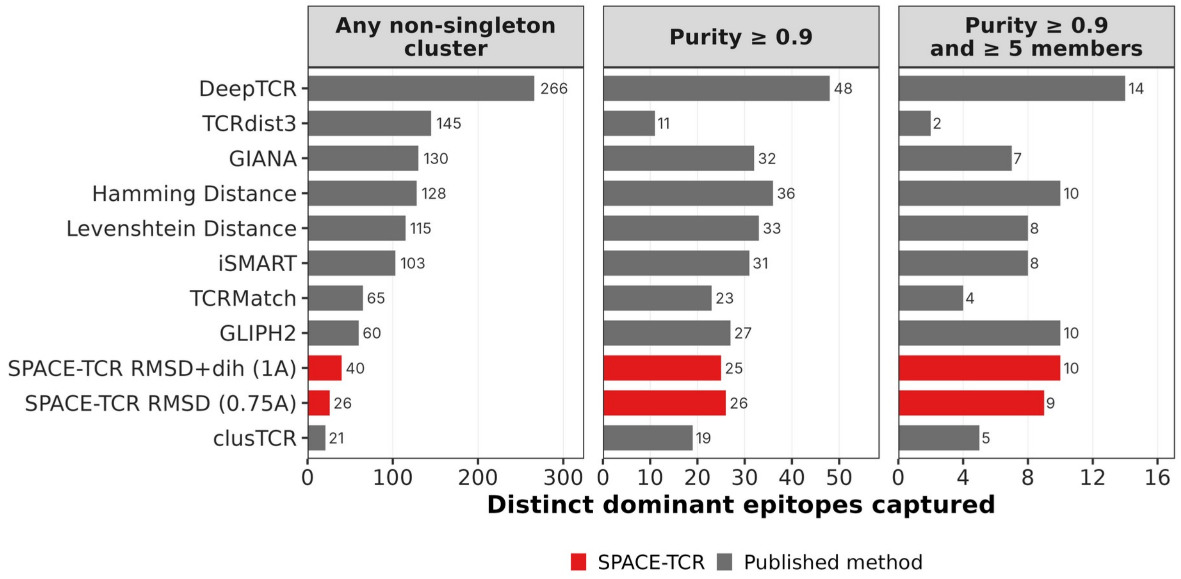


**Figure S1.** Number of dominant epitopes captured at three quality thresholds: any non-singleton cluster, purity ≥ 0.9, and purity ≥ 0.9 with at least five members. For SPACE-TCR at 0.75 Å, every epitope reached at the first threshold also appears in a high-purity cluster.

**Supplementary Table S2.** Distinct dominant epitopes captured at three cluster-quality thresholds.

| Method | Any cluster | Purity ≥ 0.9 | Purity ≥ 0.9 & size ≥ 5 | High-purity clusters | Largest cluster |
|---|---|---|---|---|---|
| SPACE-TCR (RMSD, 0.75 Å) | 26 | 26 | 9 | 141 | 56 |
| SPACE-TCR (RMSD+dih, 1.0 Å) | 40 | 25 | 10 | 131 | 107 |
| ClusTCR | 21 | 19 | 5 | 105 | 39 |
| GLIPH2 | 60 | 27 | 10 | 280 | 190 |
| TCRMatch | 65 | 23 | 4 | 172 | 14 |
| iSMART | 103 | 31 | 8 | 200 | 37 |
| Levenshtein distance | 115 | 33 | 8 | 164 | 153 |
| Hamming distance | 128 | 36 | 10 | 194 | 68 |
| GIANA | 130 | 32 | 7 | 206 | 45 |
| DeepTCR | 266 | 48 | 14 | 315 | 29 |
| TCRdist3 | 145 | 11 | 2 | 49 | 34 |

Table S2b identifies the high-purity epitopes shared by SPACE-TCR and ClusTCR. Eighteen were common to both methods; SPACE-TCR reached eight additional epitopes and ClusTCR one. Seven of the eight SPACE-TCR-only epitopes were represented by two-member clusters and RPPIFIRRL by a five-member cluster, so most remain near the detection limit. None appeared in any ClusTCR cluster, although several were recovered at high purity by other sequence methods. They therefore reflect the specific stringency of ClusTCR rather than epitopes accessible only to structural analysis. FLCMKALLL illustrates the distinction between cluster purity and Jouannet sensitivity: it dominated a high-purity SPACE-TCR cluster, but that cluster was too small to satisfy the sensitivity rule. SLLMWITQC (NY-ESO-1) was absent from the ClusTCR output.

**Supplementary Table S2b.** Epitopes dominating a high-purity cluster under focal SPACE-TCR and ClusTCR.

| Recovered by | Epitopes |
|---|---|
| Both methods (18) | AAGIGILTV, CINGVCWTV, DATYQRTRALVR, GILGFVFTL, GLCTLVAML, LLFGYPVYV, LLWNGPMAV, LLYDANYFL, LTDEMIAQY, NLVPMVATV, NQKLIANQF, NYNYLYRLF, PTDNYITTY, RPHERNGFTVL, SPRWYFYYL, TAFTIPSI, TTDPSFLGRY, YLQPRTFLL |
| SPACE-TCR only (8) | EPLPQGQLTAY, FLCMKALLL, FLRGRAYGL, ILAKFLHWL, QELIRQGTDY, RPPIFIRRL, SLLMWITQC, TPRVTGGGAM |
| ClusTCR only (1) | FTSDYYQLY |

### S3. Consensus clustering summary

Table S3 gives the values plotted in Figure S7 on the shared universe of 6,163 unique CDR3β sequences. Each consensus increased the proportion of clusters at purity ≥ 0.9 relative to both parents, while retention remained below that of the less retentive parent, as required by intersection.

**Supplementary Table S3.** Purity and retention for four base methods and four consensus settings on the shared unique-CDR3β denominator.

| Method or consensus | % clusters purity ≥ 0.9 | Global purity | Retention | Unique CDR3β clustered |
|---|---|---|---|---|
| SPACE-TCR (RMSD+dih, 1.0 Å) + GLIPH2 | 99.1% | 99.7% | 4.36% | 269 |
| SPACE-TCR (RMSD, 0.75 Å) + GLIPH2 | 99.1% | 99.6% | 3.65% | 225 |
| SPACE-TCR (RMSD, 0.75 Å) + ClusTCR | 98.4% | 99.4% | 2.21% | 136 |
| SPACE-TCR (RMSD+dih, 1.0 Å) + ClusTCR | 98.4% | 99.4% | 2.43% | 150 |
| ClusTCR | 96.3% | 98.8% | 3.26% | 201 |
| SPACE-TCR (RMSD, 0.75 Å) | 94.7% | 98.7% | 7.24% | 446 |
| SPACE-TCR (RMSD+dih, 1.0 Å) | 73.7% | 91.4% | 10.42% | 642 |
| GLIPH2 | 73.4% | 90.2% | 11.57% | 713 |

### S4. Repertoire-noise benchmark: per-subset values

Table S4 gives purity at each noise level, and Table S4b gives the corresponding mean cluster size, signal recovery, and absolute yield. At the focal 0.75 Å threshold, SPACE-TCR had the highest purity at all six noise levels. The three methods whose mean cluster size increased under dilution - ClusTCR and the Hamming and Levenshtein baselines - all use an absolute edit-distance criterion. At the highest noise level, they also had large numbers of clustered sequences and low purity, consistent with unannotated sequences being absorbed into existing clusters. SPACE-TCR and the motif methods instead decreased in mean cluster size as noise increased.

tcrdist3 behaved differently from the other methods. Its signal recovery declined from 83.1% to 36.5%, while the absolute number of clustered sequences increased to 7,533, the highest total in the benchmark. It therefore clustered most of the available universe at every noise level. This explains its low purity on the undiluted subset and shows why recovery and purity must be interpreted together.

**Supplementary Table S4.** Cluster purity at each level of the 10X donor-1 noise series, restricted to CDR3β sequences with a paired structure. DeepTCR output was unavailable for the two densest subsets.

| Method | 0% | 62.2% | 76.7% | 83.2% | 86.8% | 89.2% |
|---|---|---|---|---|---|---|
| SPACE-TCR 0.25 Å | 1.000 | 1.000 | 1.000 | 1.000 | 0.929 | 0.929 |
| SPACE-TCR 0.5 Å | 0.953 | 0.889 | 0.833 | 0.747 | 0.695 | 0.624 |
| SPACE-TCR 0.75 Å | 0.967 | 0.806 | 0.624 | 0.475 | 0.368 | 0.304 |
| SPACE-TCR 1.0 Å | 0.868 | 0.507 | 0.320 | 0.235 | 0.178 | 0.145 |
| SPACE-TCR 1.25 Å | 0.739 | 0.359 | 0.218 | 0.154 | 0.118 | 0.096 |
| iSMART | 0.947 | 0.488 | 0.282 | 0.204 | 0.154 | 0.117 |
| GIANA | 0.946 | 0.517 | 0.298 | 0.213 | 0.161 | 0.119 |
| TCRMatch | 0.940 | 0.657 | 0.377 | 0.272 | 0.209 | 0.150 |
| GLIPH2 | 0.925 | 0.601 | 0.339 | 0.240 | 0.180 | 0.140 |
| ClusTCR | 0.875 | 0.429 | 0.241 | 0.169 | 0.125 | 0.097 |
| Hamming distance | 0.875 | 0.426 | 0.235 | 0.165 | 0.123 | 0.094 |
| Levenshtein distance | 0.813 | 0.329 | 0.176 | 0.117 | 0.089 | 0.073 |
| DeepTCR | 0.718 | 0.318 | 0.207 | 0.153 | — | — |
| TCRdist3 | 0.656 | 0.325 | 0.238 | 0.173 | 0.135 | 0.111 |

**Supplementary Table S4b.** Mean cluster size, annotated-signal recovery, and absolute yield at the lowest and highest noise levels.

| Method | Mean size, 0% | Mean size, 89.2% | Recovery, 0% | Recovery, 89.2% | Sequences clustered, 89.2% |
|---|---|---|---|---|---|
| SPACE-TCR 0.5 Å | 3.56 | 2.95 | 1.8% | 2.1% | 87 |
| SPACE-TCR 0.75 Å | 3.62 | 3.13 | 3.7% | 5.1% | 475 |
| SPACE-TCR 1.0 Å | 3.78 | 3.55 | 8.4% | 17.8% | 2,897 |
| ClusTCR | 4.22 | 5.72 | 9.7% | 22.7% | 4,467 |
| Hamming distance | 4.30 | 6.03 | 9.7% | 22.8% | 4,481 |
| Levenshtein distance | 4.40 | 8.97 | 11.3% | 27.0% | 5,527 |
| iSMART | 4.47 | 3.66 | 6.4% | 17.0% | 3,109 |
| GIANA | 3.67 | 2.94 | 6.2% | 14.9% | 2,781 |
| GLIPH2 | 3.62 | 2.39 | 8.8% | 16.0% | 2,987 |
| TCRMatch | 3.19 | 2.35 | 3.5% | 5.7% | 866 |
| TCRdist3 | 3.07 | 2.60 | 83.1% | 36.5% | 7,533 |

### S5. Repertoire-noise benchmark: provenance and label reconstruction

Table S4c describes the noise series. Each subset adds 4,737 sequences, so the series is arithmetic rather than geometric and reaches 89.2% unannotated content. Restriction to CDR3β sequences with a paired structure retained 84.7-86.2% of every subset, indicating similar coverage at each noise level. The number of annotated paired sequences remained fixed at 2,374; the series therefore dilutes one annotated core rather than adding annotated and unannotated receptors together.

**Supplementary Table S4c.** Construction of the 10X donor-1 noise series and restriction to receptors with a paired structure.

| Subset | CDR3β in subset | With a paired structure | % paired | Annotated (paired) | % unannotated |
|---|---|---|---|---|---|
| 1 | 2,876 | 2,478 | 86.2% | 2,374 | 0.0% |
| 2 | 7,613 | 6,492 | 85.3% | 2,374 | 62.2% |
| 3 | 12,350 | 10,500 | 85.0% | 2,374 | 76.7% |
| 4 | 17,087 | 14,469 | 84.7% | 2,374 | 83.2% |
| 5 | 21,824 | 18,497 | 84.8% | 2,374 | 86.8% |
| 6 | 26,561 | 22,560 | 84.9% | 2,374 | 89.2% |

The original benchmark label file was not distributed, so dextramer annotations were reconstructed from the 10X binarised matrix using the published procedure: the mean dextramer score was calculated per CDR3β, a score above 10 defined binding, and all positive dextramers were retained. This produced 3,216 annotated CDR3β sequences across 39 epitopes, compared with 2,876 sequences in the original annotated subset. Of the reconstructed labels, 2,760 fell within that subset and 456 outside it; 116 of the original 2,876 were unlabelled by the reconstruction. Labels were therefore restricted to the original annotated subset so that the benchmark's definition of the signal set was preserved.

This mismatch has two small effects. The 116 original sequences not recovered by the reconstruction are treated as unannotated and therefore dilute purity equally for every method. Conversely, 51 reconstructed labels occur on sequences treated as noise in the series. A method can therefore be penalised for correctly grouping one of these receptors with a matching annotated receptor. At 51 of 22,560 sequences in the densest subset, the effect is too small to explain the reported rankings, but it makes the absolute purity values slightly conservative.

### S6. Cluster-level overview of SPACE-TCR and ClusTCR

Figure S2 presents the modelled cohort at cluster level. Each dot is a clustered TCR and each boundary denotes one cluster. Rows compare SPACE-TCR at 0.75 Å (149 clusters; 681 TCRs), SPACE-TCR at 1.0 Å (202; 1,000), and ClusTCR (111; 426). SPACE-TCR formed more clusters and clustered more receptors at both thresholds. The counts in brackets are receptors, which is what the panels draw; on the unique-CDR3β unit used for every cross-method comparison in this paper the focal figures are 446 against 201 (Table 2). Most cluster boundaries were dominated by one colour for both methods, consistent with the purity and polyspecificity results. Loosening the SPACE-TCR threshold increased the number of represented epitopes while reducing per-cluster purity, and distributed the GILGFVFTL receptors across more, smaller structural clusters.

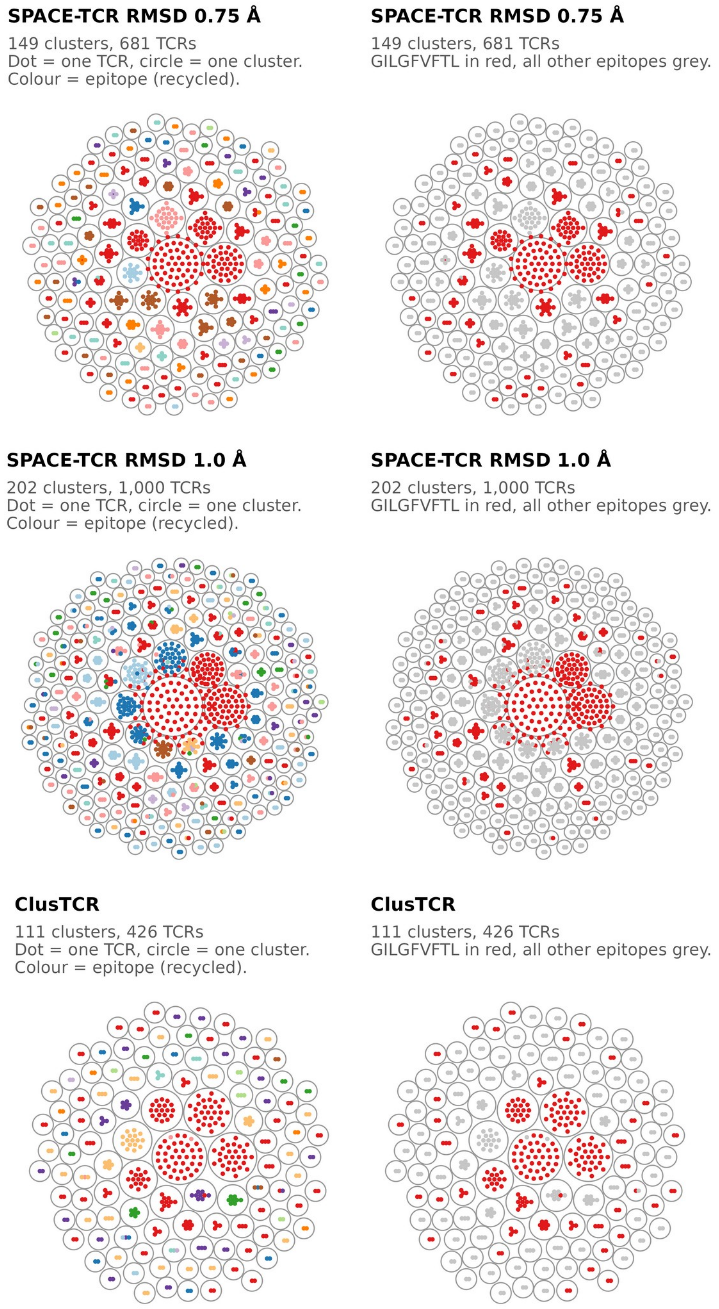


**Figure S2.** Cluster-level overview of SPACE-TCR and ClusTCR. Rows show SPACE-TCR at 0.75 Å (149 clusters; 681 receptors), SPACE-TCR at 1.0 Å (202; 1,000), and ClusTCR (111; 426). Receptor counts are given here because each dot in the panels is one clustered receptor; the unique-CDR3β figures used for every cross-method comparison in this paper are in Table 2. The right column highlights GILGFVFTL on the same cluster arrangements. Dot colour marks epitope identity and is recycled across epitopes, so it is not individually keyed; the panel's point is that cluster boundaries are internally one colour.

### S7. Behaviour of the dihedral filter across the threshold sweep

Figure S3 evaluates RMSD-only clustering and dihedral gates of <0.5, <0.3, and <0.2 radians across all five RMSD thresholds. At 0.25 Å, every setting gave 100% purity and 100% of clusters at purity ≥ 0.9, with retention of 1.28-1.36%. At 0.5 Å, global purity remained 98.8-99.0% and high-purity cluster rates 94.9-95.8%; the gate therefore added little when RMSD was already stringent. At 1.25 Å, the <0.2-radian gate increased global purity from 78.4% to 98.3% and high-purity clusters from 41.7% to 92.2%, while reducing retention from 32.5% to 7.5%. At 1.0 Å, the same gate increased purity from 92.4% to 98.8% and high-purity clusters from 75.2% to 94.5%, with retention decreasing from 20.9% to 7.0%.

The 0.5-radian gate reported in the main comparison is the least restrictive setting. At 1.25 Å, it increased purity to 84.3% and high-purity clusters to 53.8% while retaining 24.8% of the input; at 1.0 Å, the corresponding values were 92.7%, 76.2%, and 18.9%. A <0.2-radian gate largely displaced RMSD as the binding criterion: thresholds from 0.5 to 1.25 Å converged on 98.3-98.8% purity and 5.0-7.5% retention. This near-threshold-independent mode is highly selective and was therefore not used for the main benchmark. Full values are provided in sptce_dihedral_mode_metrics.csv.

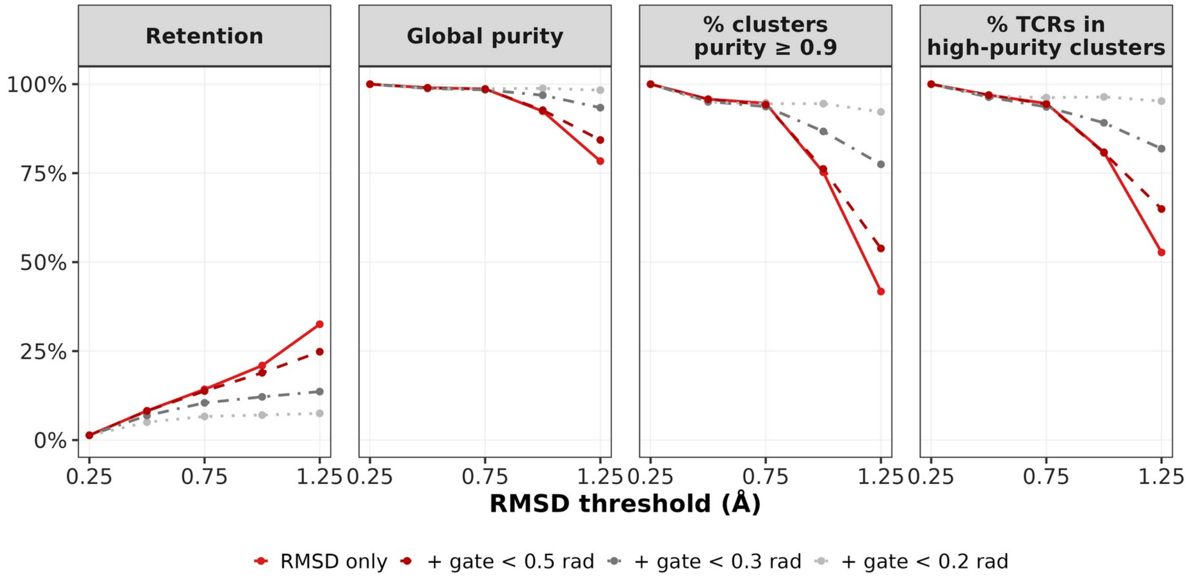


**Figure S3.** Effect of dihedral-gate stringency across five RMSD thresholds. Panels show retention, global purity, the proportion of clusters at purity ≥ 0.9, and the proportion of TCRs in high-purity clusters. Dihedral gating has little effect at tight RMSD thresholds but substantially increases purity at looser thresholds.

### S8. Composition of contamination in the GILGFVFTL clusters

Figure S4 identifies the non-GILGFVFTL receptors placed in GILGFVFTL-dominant clusters. Totals followed the overall purity ranking. SPACE-TCR at 0.75 Å admitted two non-focal receptors from two epitopes; ClusTCR admitted three from two; the 1.0 Å dihedral-gated SPACE-TCR mode admitted 12 from 10; TCRMatch 20 from 12; and GLIPH2 47 from 20. At the other extreme, tcrdist3 admitted 319 receptors from 125 epitopes, Levenshtein 252 from 91, and DeepTCR 169 from 87. The number of distinct contaminating epitopes helps distinguish repeated members of one related family from a broad scatter of unrelated labels.

Two recurrent contaminants merit attention. NLVPMVATV was the most frequent contaminant for seven of the eleven methods, including SPACE-TCR, and appeared among the top three for nine. It is also the second most abundant specificity in the cohort, so its frequency may largely reflect prevalence. The shared HLA-A*02:01 restriction with GILGFVFTL nevertheless makes a biological relationship plausible and suitable for direct testing.

GILEFVFTL, which differs from GILGFVFTL by one residue, was the leading contaminant for GLIPH2 and ClusTCR and appeared in five other methods. Grouping receptors assigned to single-substitution variants is not necessarily a biological error, but the benchmark purity metric treats the peptide strings as unrelated labels. The same issue applies to LLFGYPVYV and LLFGFPVYV. Contamination counts are therefore upper bounds on error, especially for the methods with the highest cluster purity: two of the three non-focal receptors admitted by ClusTCR were variant-peptide cases. The long tails of singleton contaminants in broader methods are less readily explained in this way.

Purity alone cannot make this distinction. Future benchmarks could collapse single-substitution peptide variants to a shared label, or report both exact-label and variant-collapsed scores. This would separate probable cross-recognition from unrelated contamination.

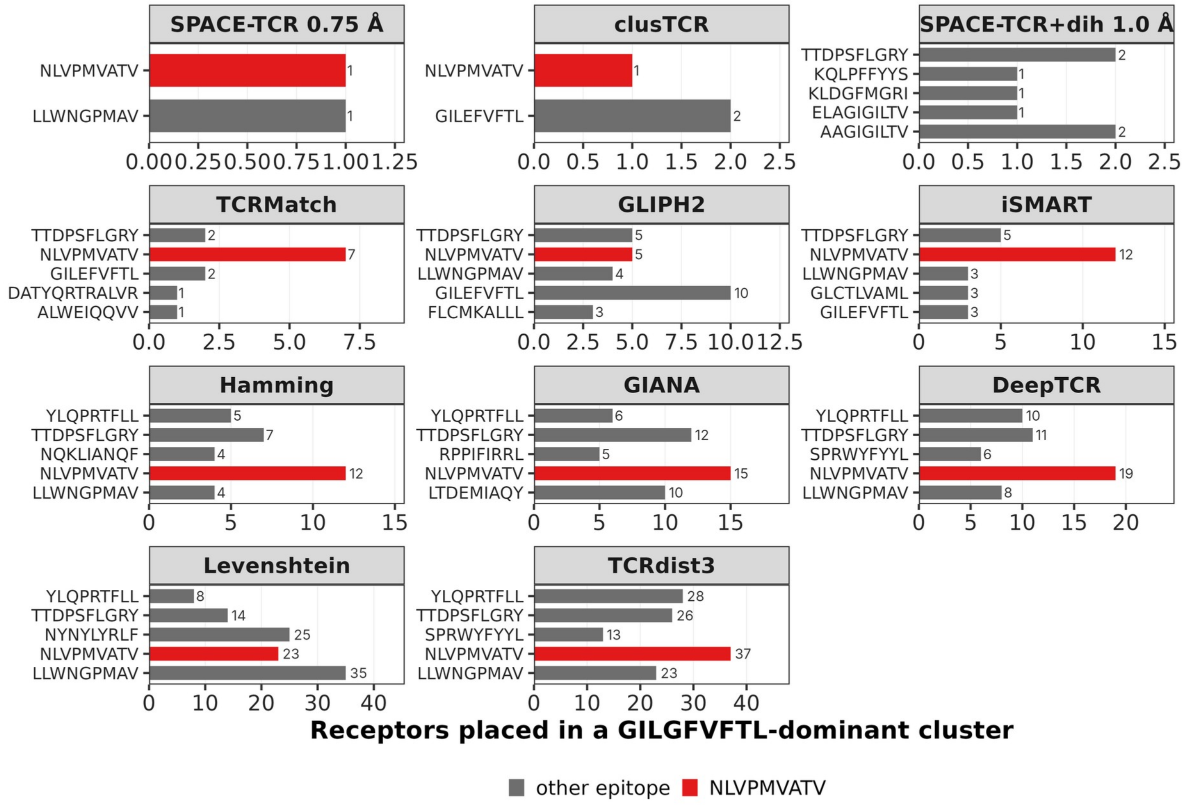


Five most frequent contaminating labels per method; note the free x scale.

**Figure S4.** Non-GILGFVFTL labels within GILGFVFTL-dominant clusters. NLVPMVATV is highlighted. Focal SPACE-TCR admitted two non-focal receptors and ClusTCR three, compared with 319 receptors across 125 epitopes for tcrdist3. GILEFVFTL, a single-substitution peptide variant, was the leading contaminant for GLIPH2 and ClusTCR.

### S9. Data and code availability

All associated data and code are available (https://github.com/dstcr/SPACE-TCR-repo). Per-cluster membership tables for every method and threshold are provided as supplementary data, including cluster identifiers, member TCR identifiers, paired CDR3α and CDR3β sequences, V- and J-gene calls, epitope labels, cluster purity, and cluster size. The predicted paired structures are also provided - https://zenodo.org/records/23765886 . The deposited repository additionally contains both benchmarks' input tables and the structural output they were clustered into, the pairwise CDR-loop distance tables and the CDR3 dihedral tables for the curated cohort, together with the verification script, so the reported clustering can be re-derived at any threshold at or below 1.25 Å without repeating structure prediction. The sequence-only distillation code and its complete intermediate results are deposited with it (Supplementary Section S12).

### S10. Complete repertoire-noise benchmark output

This section provides a long-format listing of the complete supplied repertoire-noise benchmark, including all five SPACE-TCR RMSD thresholds, RMSD-only and dihedral-gated settings, and the published comparison methods. Purity, mean cluster size, cluster count, and the proportion of each published method's clustered output retained after restriction to paired receptors are reported as produced by the benchmark run. DeepTCR output was unavailable for subsets 5 and 6. The exact console output and R-readable CSV files are supplied with the supplementary data. The composition of the six subsets after restriction to receptors with a paired structure is given once, in Supplementary Table S4c.

Note. The number of labelled paired CDR3β sequences is fixed at 2,374 across all subsets.

### S11. Reproducing the benchmark from the deposited data

The repository contains the input tables and structural measurements used for clustering in both benchmarks, so every number in Tables 1 and 2 is re-derivable without rerunning structure prediction. For the curated cohort this is the 4,779-receptor input table, 828 gzipped pairwise distance tables and the CDR3 dihedral table (5.9 MB in total); for the noise benchmark it is the 23,529-receptor table of the largest subset with a membership column giving each receptor the first subset in which it appears, together with the reference subset's distance tables and dihedrals (30 MB). The smaller subsets are strict restrictions of the largest by receptor identifier, with identical epitope labels throughout, so they are derived rather than stored.

A verification script reconstructs clusters as connected components at each distance threshold, recalculates purity for clusters of at least two receptors, and compares the results with the deposited tables. All 36 assertions pass. The focal operating point reproduces exactly: 149 clusters, 681 receptors clustered, 94.63% of clusters at purity ≥ 0.9, and a pooled purity of 0.9868 against the 98.7% reported in Section 3.1. All six per-epitope sensitivity denominators reproduce exactly against the input table. The script validates the analysis layer and the integrity of the deposited coordinates; it does not re-predict any structure, and the structural step is evidenced separately by the training-overlap analysis of Section 3.8.

### S12. Sequence-only distillation of the structural clustering

Structure prediction is the throughput constraint on this method, so we tested whether the clustering could be distilled into a sequence-only model that would propose candidate groups for structural modelling rather than requiring every receptor to be folded. A siamese encoder was trained on SPACE-TCR's own cluster assignments, embedding each receptor from its CDR3β and CDR3α sequences and its TRBV and TRBJ identities with no structural input: a 24-dimensional learned amino-acid embedding, convolutional kernels of width 3, 5 and 7, and an L2-normalised 64-dimensional output trained under a contrastive loss with margin 0.7, where a pair is positive when both receptors occupy the same structural cluster. Clusters were formed from the embedding by the same connected-components procedure used throughout, swept across distance thresholds.

Evaluation held out whole epitopes rather than individual receptors, because two receptors for the same epitope are frequently near-clonal and a receptor-level split allows a model to memorise the epitope without generalising. Thirty per cent of epitopes were withheld, test receptors sharing a CDR3β with any training receptor were discarded, and scoring covered every cohort receptor carrying a held-out epitope whether or not SPACE-TCR had clustered it, so receptors the structural method missed counted against it. Weight selection additionally required a minimum embedding spread: a contrastive encoder can collapse to a single direction and still rank pairs correctly, because the distances remain ordered while shrinking towards zero, and such an encoder assigns every receptor to one component at any threshold.

The sequence-only model did not reliably reproduce the structural clustering. We identified three limitations. First, the encoder reproduces the clustering only when supervised by the permissive 1.25 Å assignment; at the focal 0.75 Å threshold it does not, and the 1.25 Å result is an operating-point artefact rather than a property of the model. Second, it does not transfer: applied unchanged to the repertoire-noise benchmark it absorbs the unannotated background instead of rejecting it, which is the specific failure the structural method avoids. Third, restricting the teacher to high-purity clusters, which appears to strengthen the supervision, leaks the epitope label into it, since a cluster retained because its members share an epitope makes "same cluster" approach "same epitope"; inner-validation AUROC reached 1.000, which is not a credible generalisation score on a training set of this size. That filter is also unusable in practice, because it requires epitope labels for the whole repertoire, and a fully labelled repertoire needs no clustering method.

Structure prediction therefore remains the principal constraint on throughput. The tested sequence-only model did not reliably reproduce the structural clustering. Further evaluation on larger paired TCR datasets is needed to determine whether this limitation extends beyond the present model and training set. The code and the full set of intermediate results are deposited so that the experiment can be repeated on a larger cohort.

### S13. The unit of comparison across published methods

The nine published methods are not all measured in the same unit, and retention figures should be read with that in mind. Three unit classes arise in the benchmark: the paired receptor, which is the unit SPACE-TCR clusters; the receptor keyed on its β chain, which is the unit of the methods operating on CDR3β with its gene calls; and a pooled single-chain list in which α and β sequences are mixed into one input, which is the unit of the methods supplied with that list by the reference benchmark. For the cohort used here these give 4,779, 4,292 and 8,395 units respectively, and the per-epitope denominators differ correspondingly — for GILGFVFTL, 705, 455 and 938. These per-epitope denominators count a sequence under every epitope it is labelled with; see Supplementary Section S1 for the thirteen sequences this affects. A method scored over a larger universe is not thereby worse, and the three classes should not be ranked against one another on retention without restatement in a common unit. Purity is less affected, because it is a within-cluster quantity, but it is not wholly immune. Section 3.2's head-to-head comparison restricts all methods to the shared unique-CDR3β universe for this reason.

### S14. Structural neighbour availability and what retention measures

Retention in this benchmark is dominated by how many receptors have any structural neighbour at all, rather than by how the remainder are partitioned. Of the 4,779 receptors, 1,555 have at least one neighbour within 1.25 Å, the most permissive threshold evaluated; the other 3,224, including the 748 receptors without a structural model, are singletons at every threshold and cannot be clustered by any setting of this method. 9,642 receptor pairs fall within 1.25 Å in total. The retention column should therefore be read as the product of two things: the fraction of the repertoire with a structural neighbour, which is a property of the cohort and of the predictor's resolution, and the threshold's selectivity among those that have one. A method clustering on CDR3β edit distance faces no equivalent floor, which is part of why the sequence methods retain more, and is a further reason to compare the two families at matched purity rather than on retention alone.

### S15. Paired-chain convergence: distinct αβ pairings within a single epitope

Figure S5 supports Section 3.6 of the main text.

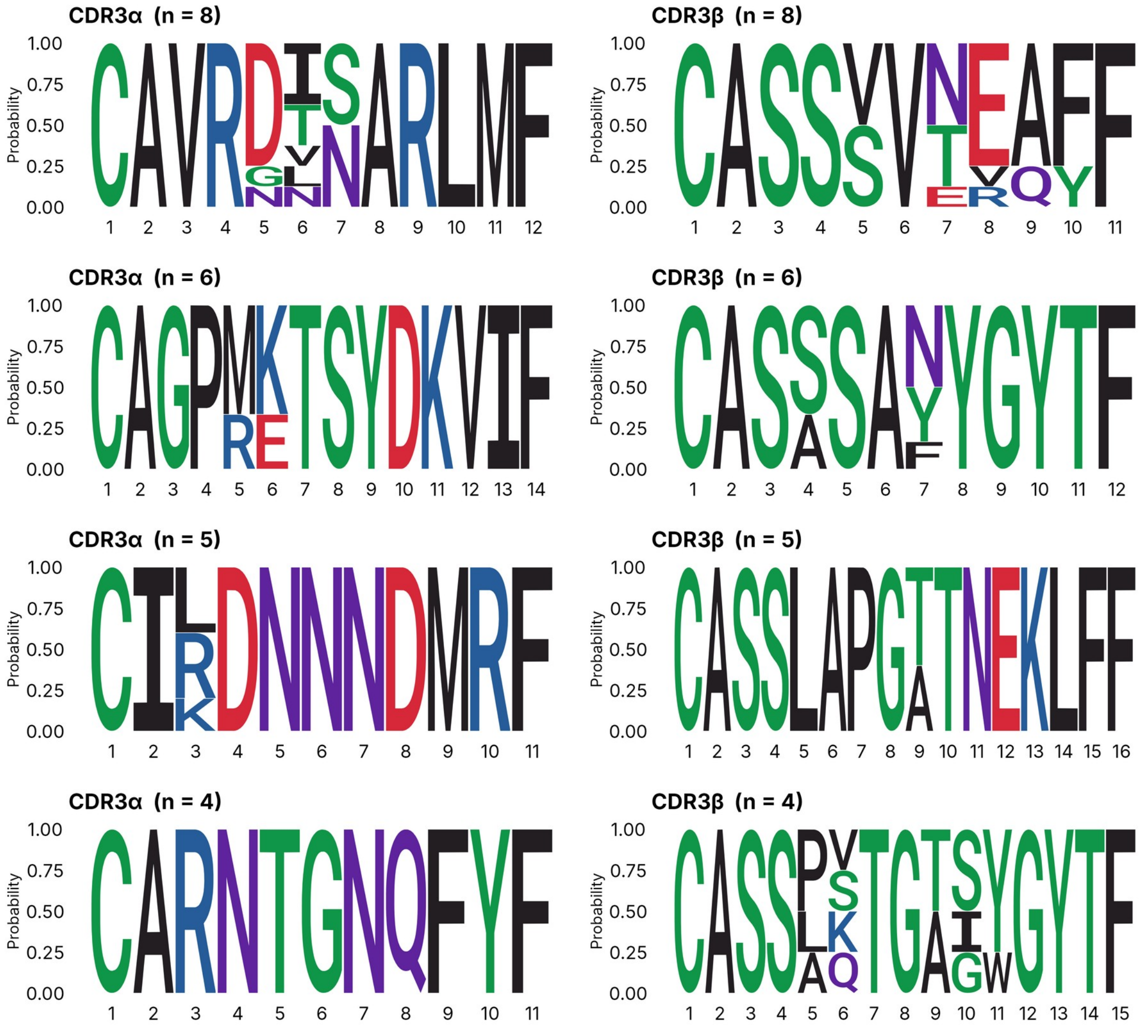


Four largest high-purity NLVPMVATV clusters at the focal threshold; one cluster per row, paired α (left) and β (right). R

**Figure S5.** Paired CDR3α and CDR3β sequence logos for high-purity NLVPMVATV clusters at the focal threshold. Each row represents one cluster and shows a recurrent α motif paired with a specific β motif, resolving the epitope into distinct paired clonotype families.

### S16. Training-data leakage and held-out validation

Figure S6 supports Section 3.8 of the main text.

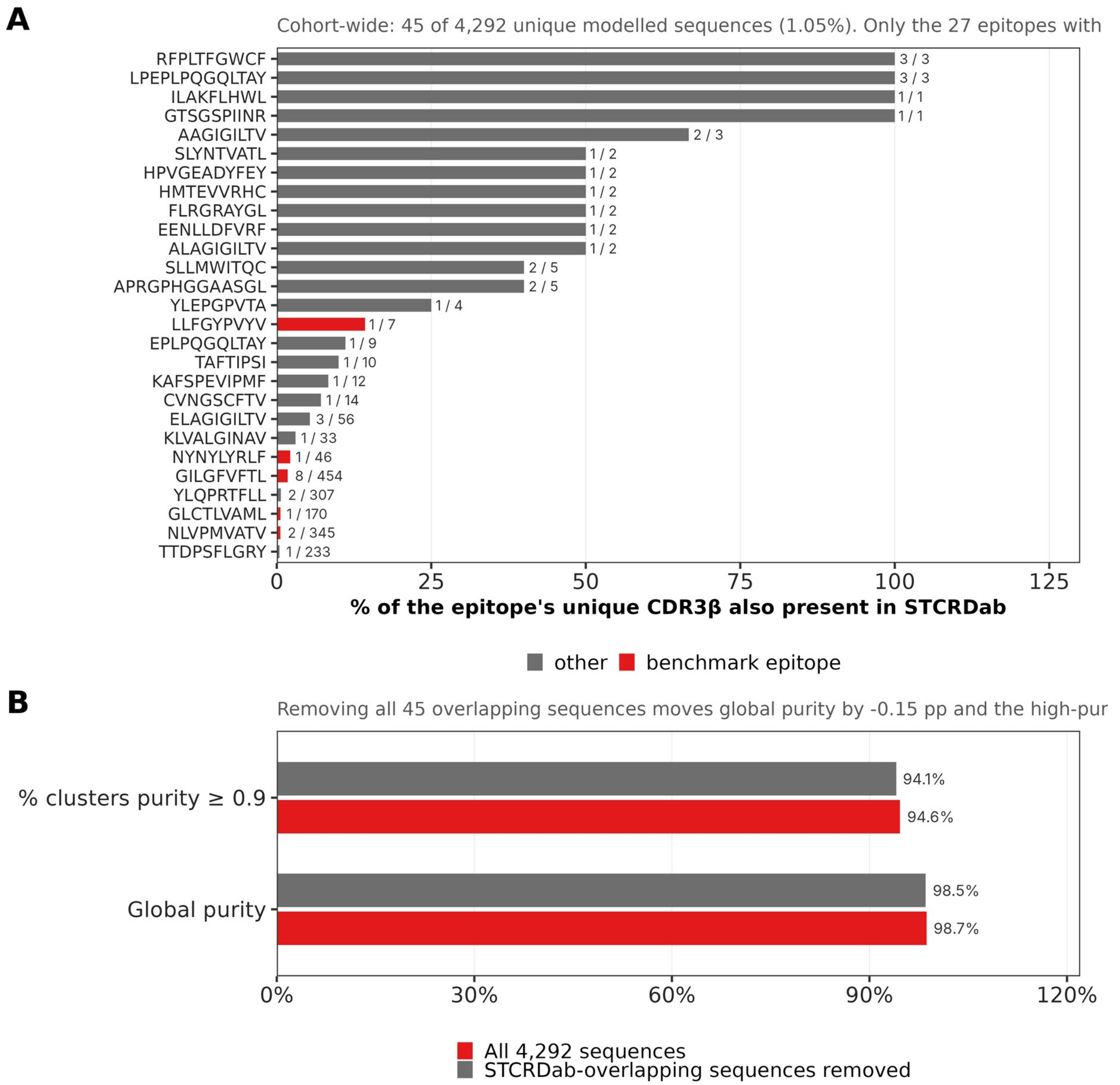


**Figure S6.** Training-set overlap and held-out validation. (A) Overlap between cohort CDR3β sequences and STCRDab was 45 of the cohort's 4,292 unique CDR3β sequences (1.05%). Per-epitope overlap is reported over each epitope's own unique CDR3β count, with each of the thirteen two-label sequences assigned to a single epitope (Supplementary Section S1), and is therefore dominated by the rare epitopes: four epitopes represented by one or three sequences are wholly contained in STCRDab, while among the 47 epitopes with at least ten unique CDR3β the maximum is 10.0% and the median 0%. For the six benchmark epitopes the overlap is 8 of 454 for GILGFVFTL (1.8%), 2 of 345 for NLVPMVATV, 1 of 170 for GLCTLVAML, 1 of 46 for NYNYLYRLF, 0 of 81 for FLCMKALLL, and 1 of 7 for the sparse LLFGYPVYV. (B) Removing all overlapping CDR3β sequences changed the focal purity metrics by less than one percentage point.

### S17. Consensus clustering with sequence-based methods

Figure S7 supports Section 3.10 of the main text.

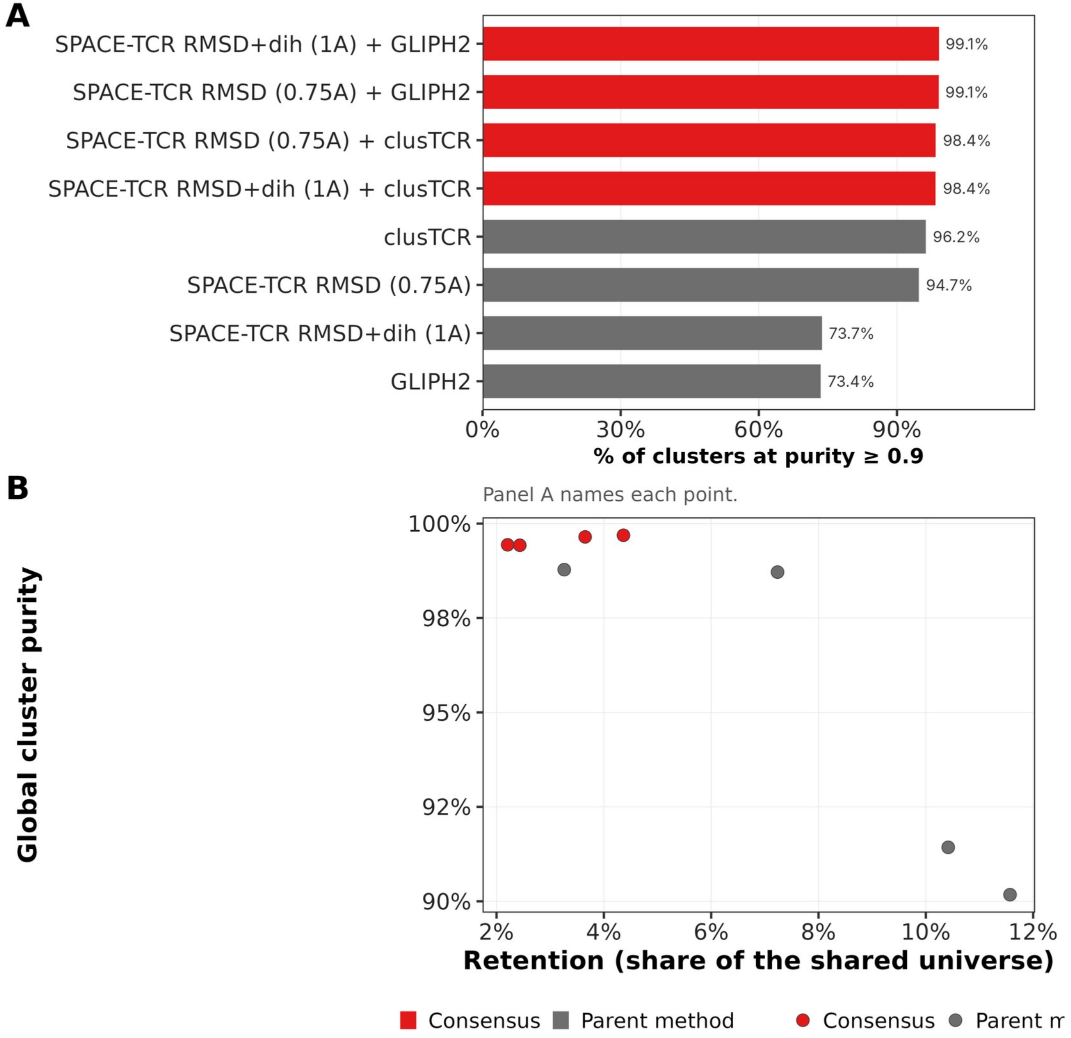


**Figure S7.** Consensus clustering on a shared universe of 6,163 unique CDR3β sequences. (A) Every SPACE-TCR × sequence-method consensus exceeds both parents in the percentage of clusters at purity ≥ 0.9. (B) Consensus settings reach 99.4-99.7% global purity at 2.2-4.4% retention, illustrating the expected exchange of yield for confidence.

### S18. Formal definitions of the evaluation metrics

These are the definitions summarised in Section 2.4. Metric definitions followed the Jouannet implementation. For a clustering that partitions the input TCRs into clusters C = {$c_1$, $c_2$, ..., cK}, let |c| be the size of cluster c; let nc,e = |{t ∈ c : epitope(t) = e}| be the number of TCRs in c labelled with epitope e; and let d(c) = maxe∈E nc,e be the number carrying the dominant epitope. Non-singleton clusters are denoted $C_2$ = {c ∈ C : |c| ≥ 2}. Global purity is the fraction of all clustered TCRs that carry their own cluster's dominant epitope (Eq. S1); summing over clusters before dividing is what makes it an average over receptors rather than over clusters. Retention is the fraction of all N input TCRs assigned to non-singleton clusters (Eq. S2). The methods take different inputs — paired receptors, CDR3β with its gene calls, or a pooled single-chain list — so for every cross-method comparison N is the cohort's 4,292 unique CDR3β and the unit counted is the unique sequence. Global purity = ( Σc∈$C_2$ d(c) ) / ( Σc∈$C_2$ |c| )(S1)

= ( number of clustered TCRs carrying their cluster's dominant epitope ) / ( total number of clustered TCRs )

$$\text{Retention} = \left( \Sigma_{c\in C_2} |c| \right) / N \qquad \text{(S2)}$$

= ( number of input TCRs assigned to non-singleton clusters ) / ( total number of input TCRs )

Epitope-level sensitivity uses a stricter qualifying rule. For epitope e, let Te = {t : epitope(t) = e} be all input TCRs labelled e. In the Jouannet implementation, a qualifying e-specific cluster contains at least four TCRs, is dominated by epitope e, and has at least 67% of its members labelled e (Eq. S3). Sensitivity is the fraction of all epitope-e TCRs recovered in such clusters (Eq. S4). The numerator counts distinct receptors: a method that assigns one receptor to several qualifying clusters must not count it more than once, which would otherwise allow a value above 1. The unit of Te is the paired receptor for SPACE-TCR and the unique CDR3β sequence for the CDR3β-keyed methods, so epitope totals differ between the two groups and are given in Supplementary Section S13.

$$C_e = \{ c \in C : |c| > 3,\ n_{c,e} = d(c),\ n_{c,e}/|c| \geq 0.67 \} \qquad \text{(S3)}$$

= { clusters with at least four TCRs, dominated by epitope e, and at least 67% labelled with epitope e }

$$\text{Sensitivity}(e) = \left( \Sigma_{c\in C_e} n_{c,e} \right) / |T_e| \qquad \text{(S4)}$$

= ( number of epitope-e TCRs found in qualifying e-specific clusters ) / ( total number of epitope-e TCRs in the input data )

Polyspecificity was defined as the fraction of non-singleton clusters containing more than one epitope label. Let m(c) be the number of distinct epitopes in cluster c (Eq. S5), and let Ipoly(c) be one when m(c) > 1 and zero otherwise (Eq. S6). The global score is the mean of this indicator across non-singleton clusters (Eq. S7). Polyspecificity is stricter than purity: a cluster that is 95% one epitope and 5% another remains high-purity but is counted as polyspecific.

$$m(c) = |\{ e \in E : n_{c,e} > 0 \}| \qquad \text{(S5)}$$

= number of different epitope labels present in cluster c

$$I_{poly}(c) = \mathbb{1}\{ m(c) > 1 \} \qquad \text{(S6)}$$

= 1 if cluster c contains more than one epitope label, 0 if cluster c contains only one epitope label

$$\text{Polyspecificity} = \left( \Sigma_{c \in C_2} \mathbb{1}\{ m(c) > 1 \} \right) / |C_2| \quad \text{(S7)}$$

= ( number of non-singleton clusters containing more than one epitope label ) / ( total number of non-singleton clusters )

The two cluster-level proportions reported in Tables 1 and 2 and in Figures 1B and 2 derive from the purity of a single cluster, which Eqs S1–S7 do not define. Per-cluster purity is the dominant-epitope fraction within one cluster (Eq. S8). A cluster is high-purity when that fraction is at least 0.9, and the two reported proportions are the share of non-singleton clusters that are high-purity (Eq. S9) and the share of clustered TCRs lying in them (Eq. S10). Eq. S1 averages over receptors and Eq. S9 averages over clusters, so a cluster's influence on the first grows with its size but is fixed in the second. The two therefore separate whenever the impure clusters are not of typical size: at the focal threshold the impure ones are small, which holds Eq. S9 to 94.6% while Eq. S1 stands at 98.7%.

$$p(c) = d(c) / |c| \quad \text{(S8)}$$

= ( number of TCRs in cluster c carrying its dominant epitope ) / ( size of cluster c )

$$\text{Clusters at purity} \geq 0.9 = \left( \Sigma_{c \in C_2} \mathbb{1}\{ p(c) \geq 0.9 \} \right) / |C_2| \quad \text{(S9)}$$

= ( number of non-singleton clusters whose per-cluster purity is at least 0.9 ) / ( total number of non-singleton clusters )

$$\text{TCRs in high-purity clusters} = \left( \Sigma_{c \in C_2} |c| \cdot \mathbb{1}\{ p(c) \geq 0.9 \} \right) / \left( \Sigma_{c \in C_2} |c| \right) \quad \text{(S10)}$$

= ( number of clustered TCRs in high-purity clusters ) / ( total number of clustered TCRs )

### S19. Re-validation against functionally screened epitope labels

Figure S8 supports Section 3.11 of the main text. Labels screened by Messemaker et al. (Immunity 2026). In A and B, SPACE-TCR clusters twice as many receptors as ClusTCR at equal purity, and purity holds when non-validating receptors are relabelled to a class of their own. C and D show the share of clustered receptors whose labels were validated experimentally, with receptors split by their number of close sequence neighbours to stratify by clustering difficulty. Faded bars denote non-significant differences.

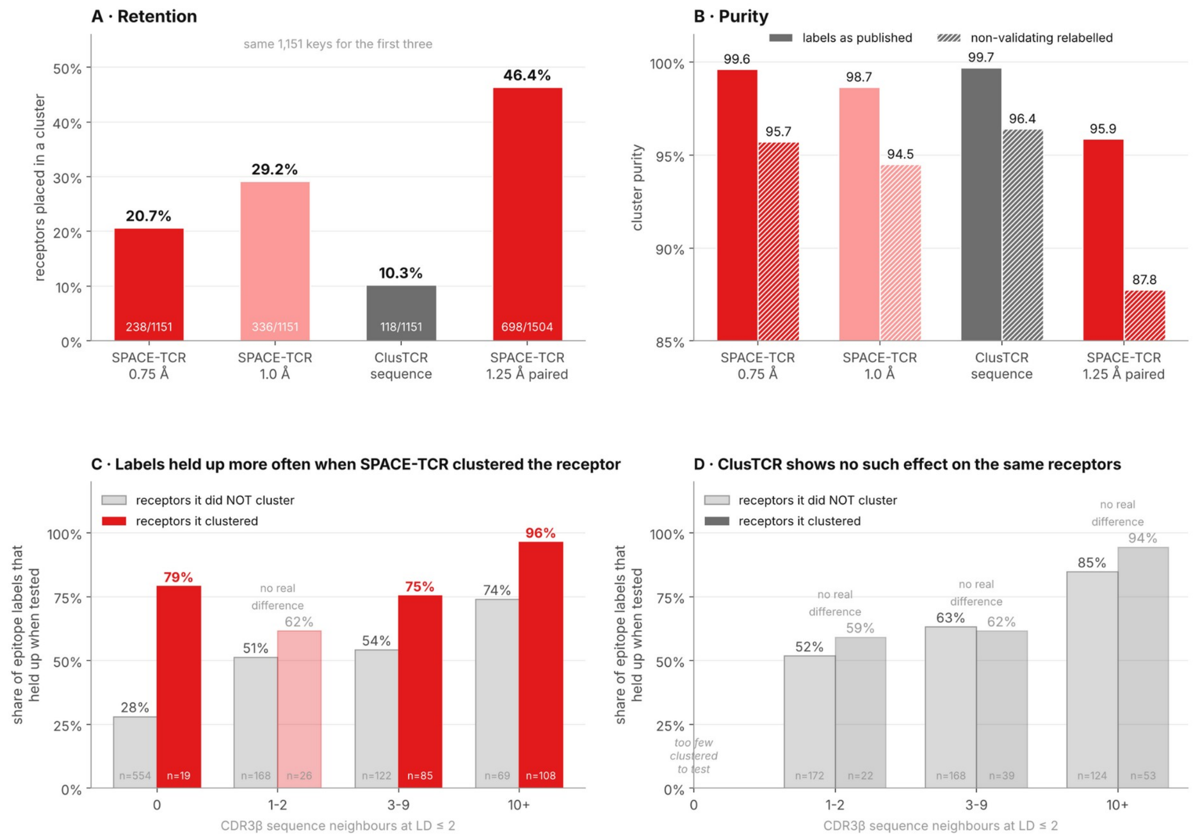


Purity axis starts at 85%. One-sided Fisher, uncorrected; SPACE-TCR's significant strata survive Bonferroni (p = 9.0e-6, 1.4e-3, 1.6e-5).

**Figure S8**. Re-validation against functionally screened epitope labels. Labels screened by Messemaker et al. (Immunity 59:2775–2788, 2026). (A) Retention, the fraction of receptors placed in a cluster, with counts inset; the first three settings share an identical 1,151-key universe. (B) Cluster purity with labels as published (solid) and with every non-validating receptor reassigned to its own class (hatched); the axis starts at 85%. (C, D) Share of epitope labels that held up under functional screening, among the receptors each method clustered versus those it left unclustered, split by the number of CDR3β neighbours at Levenshtein distance ≤ 2 so that neither method is rewarded for clustering public sequences. Faded bars mark strata where the difference is not significant (one-sided Fisher, uncorrected).

### S20. Contribution of the α chain to predicted β geometry

Figure S9 supports Section 3.6 of the main text.

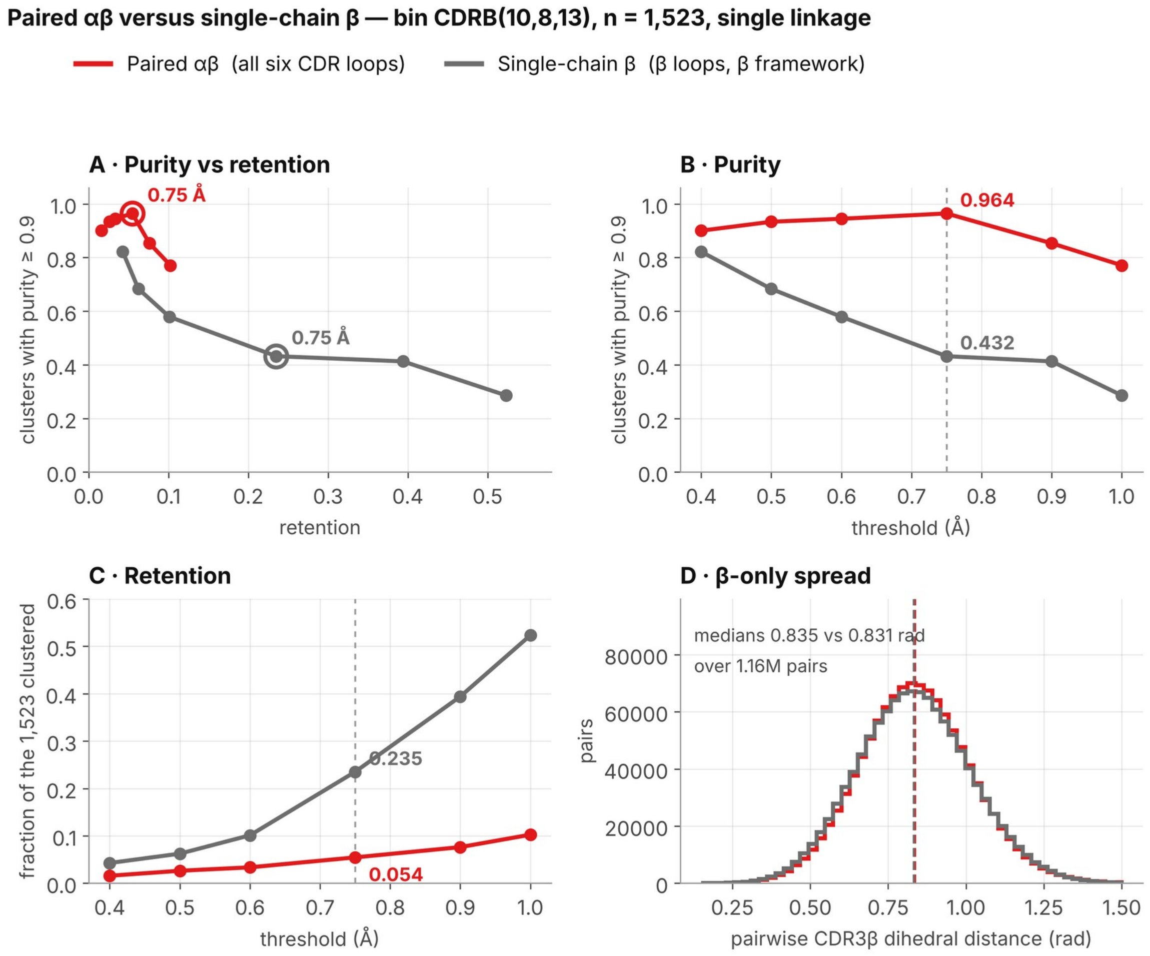


**Figure S9**. Paired αβ versus single-chain β clustering. Bin CDRB(10,8,13), n = 1,523 receptors re-modelled with a single constant α partner, so that any remaining difference between β conformations derives from the β sequence alone. Thresholds are shown over the range used in this paper, 0.4–1.0 Å, with the focal 0.75 Å point ringed in (A) and marked in (B) and (C). (A) Purity against retention. Across every comparable operating point in this range the paired method sits above the single-chain readout. (B, C) Purity and retention against threshold; each arm is scored on its own native geometry, the paired arm over all six CDR loops and the single-chain arm over β loops anchored on the β framework only, which uses no α-chain coordinates, so the thresholds are not on a common scale. (D) Pairwise CDR3β backbone dihedral distance, an internal coordinate and therefore chain-separable: the spread is unchanged between real-α and constant-α models (medians 0.835 and 0.831 rad over 1.16 million pairs), so the constant partner did not homogenise the β conformations. Single linkage throughout. A constant α is an approximation to a single-chain predictor, which does not exist in this toolchain.
